# Neural networks as decision trees: an analytical solution for learning and neural selectivity

**DOI:** 10.64898/2026.09.23.753718

**Authors:** Hugo Tissot, Jonas Ranft, Yves Boubenec

## Abstract

Nonlinear neural networks develop structured internal representations, yet how their geometry is determined by the tasks being learned remains poorly understood. Here, we develop an analytical framework for piecewise-linear feedforward and recurrent networks that links learning, activation-region structure, and neural selectivity. We show that, at any stationary point of gradient-aligned learning, a nonlinear network decomposes into local linear regressions over its activation regions. Deviations from the corresponding least-squares solutions are jointly constrained by network weights shared across regions and vanish in the low-error regime, yielding an approximate piecewise least-squares decomposition of the task. This structure admits a decision-tree interpretation, which we recover numerically by fitting trees to predict network activation patterns from the input. We further derive how the regions in which neurons are active determine the task statistics they capture, thereby organizing neural selectivity into distinct subpopulations. The resulting predictions closely match the selectivity geometry observed in simulated networks and two empirical neural datasets. Finally, we show that neural baseline regulates activation-pattern diversity, placing networks along a continuum between coarse, generalizing representations and fine-grained, expressive representations. Together, these results establish activation regions as a unifying framework for describing how nonlinear networks decompose task structure into local computations and how those computations can be revealed through neural activity.

## 1 Introduction

A central question at the interface of machine learning theory and systems neuroscience is how the observable properties of neural activity, such as tuning curves, mixed selectivity, or the geometry of population responses, are determined by the function computed by a network. For linear networks, this question has a clean answer: deep linear networks under gradient-aligned learning dynamics converge to least-squares solutions whose representational geometry is governed by the input-output covariance structure of the task [1, 2]. Both biological and artificial neural systems, however, are fundamentally *nonlinear*. Modern architectures rely extensively on activation gating and recurrent interactions, while cortical circuits exhibit strongly nonlinear response properties. Extending the linear theory to this nonlinear setting is the central theoretical challenge addressed in this work.

Piecewise-linear networks, including ReLU architectures, partition the input space into activation regions on which the network implements an affine map [3, 4]. Each region is defined by the configuration of active and inactive hidden units, and inputs that share an activation pattern share the same effective linear computation. Previous work has characterized the combinatorial and geometric properties of these regions, including their number, expressivity, and partition structure of the input space [5–8], but what these regions compute, and how learning shapes that computation, has remained largely unaddressed. In particular, how *regional* computations combine to organize the *global* geometry of neural representations is unknown.

Here, we develop an analytical framework whose ultimate goal is to understand representation geometry resulting from learning in nonlinear neural networks with piecewise-affine activations. We first show that, at any stationary point of gradient-aligned learning, nonlinear networks decompose into local linear regressions over their activation regions. We then characterize how deviations from the corresponding least-squares solutions are constrained by the network weights shared across regions. In the low-error regime, these deviations vanish, implying that expert nonlinear networks implement an approximate piecewise least-squares decomposition of the task. We derive this result for deep feedforward networks and extend it to recurrent networks at equilibrium. This framework yields a two-part functional interpretation of nonlinear computation: the routing of inputs to specific neural ensembles and the implementation of a region-specific linear regression by each ensemble. This decomposition provides a direct link between the activation structure and the geometry of neural selectivity in real neural data.

A key prediction of this framework is therefore a closed-form solution for the geometry of neural selectivity, yielding directly testable relationships between task statistics and neural population structure. The theory predicts that activation regions induce a modular organization of selectivity across neural subpopulations: neurons that share activation patterns develop similar selectivity because they implement the same local input-output mappings, whereas neurons associated with distinct regions specialize toward different task statistics. Such modular and task-dependent organization is a recurring feature of biological and artificial neural representations, where neurons often segregate into functionally specialized subpopulations with distinct selectivity geometries [9–15]. Our framework offers a mechanistic account for the emergence of this structure, with explicit analytical expressions linking the selectivity covariance of each activation-defined neural subpopulation to task statistics. We validate these predictions in trained artificial neural networks and two independent mouse prefrontal and visual cortex datasets [12, 16], finding close quantitative agreement between predicted and observed selectivity geometries.

## 2 Deep feedforward network

### 2.1 Setup

We consider an *L*-layer network with input dimension *n*_*x*_, hidden widths *m*_1_, …, *m*_*L*−1_, and output dimension *n*_*y*_. For compactness, set *m*_0_ := *n*_*x*_ and *m*_*L*_ := *n*_*y*_. The network receives a dataset of *p* input–target pairs 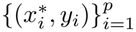, where 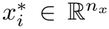 is the ordinary input of sample *i* and 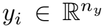 is its target. The index *i* labels samples throughout the derivation. Because the activation functions are piecewise affine, each input selects one affine piece at every hidden layer. Let 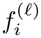 denote the piece selected by input *i* at layer *l*. The corresponding activation region is

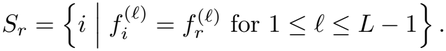

Thus, inputs belong to the same region when they select the same affine piece at every layer.

Within region *r*, the slopes and intercepts of these pieces are fixed. Since the activation function is applied independently to each unit, the slopes of the selected affine pieces at layer *l* can be collected into a diagonal matrix 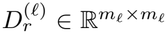. The activation function for layer *l* can then be written explicitly within region *r* as the affine transformation

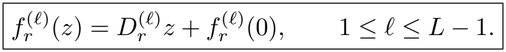

Here 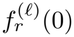 denotes the intercept of the selected affine piece. For ReLU, 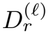 is a diagonal binary mask and 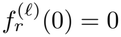. The network is therefore a composition of fixed affine maps within each activation region.

To incorporate the trainable biases into the same matrix expressions as the linear weights, we index the constant coordinate by 0 and write

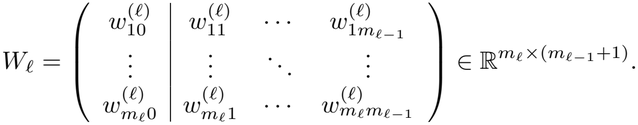

We use the compact block notation

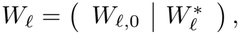

where 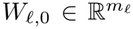 is the trainable bias vector and 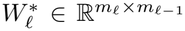 contains the weights connecting layer *l* −1 to layer *l*. The augmented input and activity vectors are ordered with their constant coordinate first.

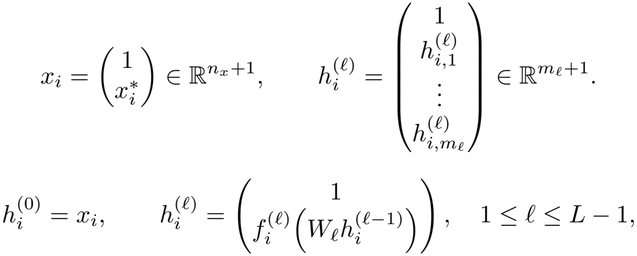

The forward pass is

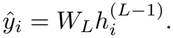

### 2.2 Affine form of the network

Within region *r*, the network is affine.

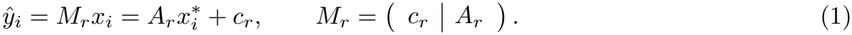

#### Linear part

Composing the regional slopes and weight matrices gives

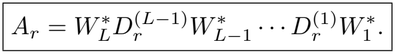

#### Offset term

Let 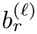 denote the regional activity offset at layer *l*. It is the activity produced by the regional affine map when the network input is zero. Starting from

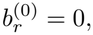

these offsets obey the recursion

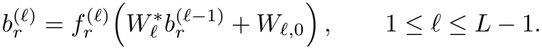

The regional output intercept is therefore

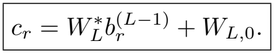

The recursion makes explicit how constant terms introduced at early layers are transformed by every subsequent layer.

The pair (*A*_*r*_, *c*_*r*_) fully characterizes the affine map in region *r*: *A*_*r*_ is its linear transformation, whereas *c*_*r*_ is its output offset.

### 2.3 Gradient with respect to *W*_*l*_

We now compute the output differential induced by a perturbation *dW*_*l*_ of the weight matrix at layer *l*, within a fixed input region *r*. Define the downstream propagation operator

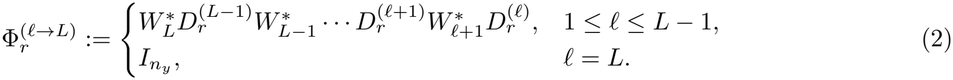

The chain rule gives the output differential

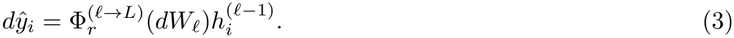

This is the standard backpropagation differential. The operator 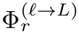 propagates the perturbation from layer *l* to the output, while 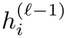 is the presynaptic activity presented to *W*_*l*_. Define the per-sample error 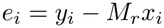. The differential of the mean-squared loss 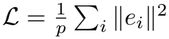 is

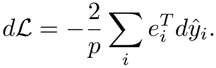

Substituting Eq. (3) and using the trace identity to isolate *dW*_*l*_ yields

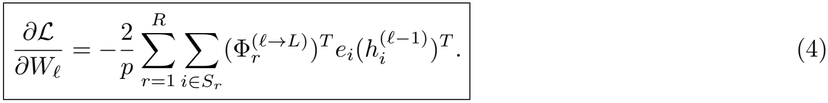

The gradient therefore decomposes into region-wise contributions. Each contribution is the sum across samples in that region of outer products between the upstream error signal and the activity entering the layer. Its first column corresponds to the gradient with respect to the trainable bias *W*_*l*,0_, while its remaining columns correspond to the gradient with respect to the weight matrix *W* ^*^.

### 2.4 Stationarity condition

For 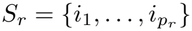 define

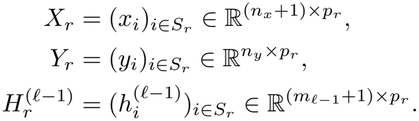

Define the augmented upstream propagation operator

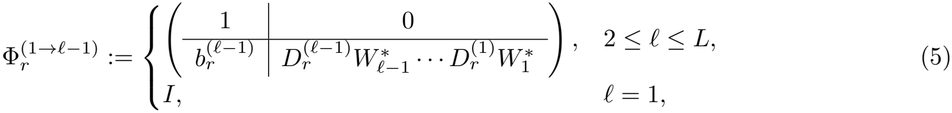

Then

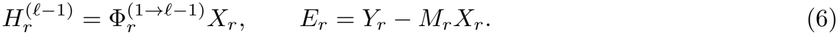

Stationarity with respect to *W*_*l*_ gives

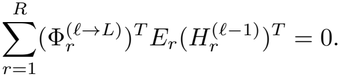

Using Eq. (6), this becomes

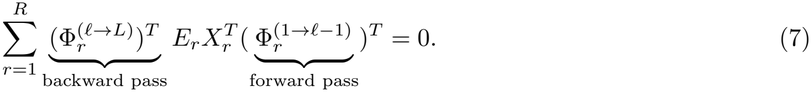

Each term in the sum is the contribution of region *r* to the gradient of *W*_*l*_. The central factor 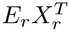 measures the correlation between prediction errors and inputs within that region. The transposed downstream operator on the left propagates these errors backward from the output to layer *l*, while the upstream operator appearing transposed on the right maps the inputs to the activities entering that layer. Because *W*_*l*_ is shared across regions, stationarity requires only that these regional gradient contributions cancel across the sum, not that each contribution vanish independently.

### 2.5 Least-squares decomposition

Assuming *X*_*r*_ has full row rank, define the regional affine least-squares estimator

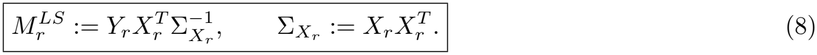

Since

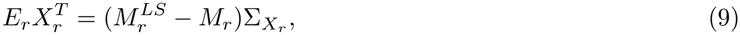

substitution into Eq. (7) gives

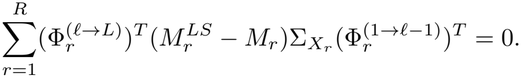

Define the regional deviation as the difference between the transformation implemented by the network and the local least-squares solution,

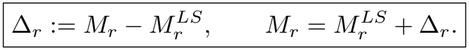

Then

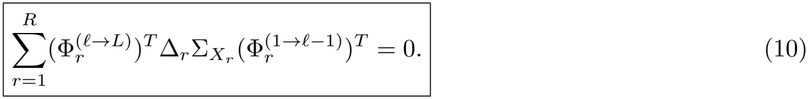

Thus, Δ_*r*_ measures how far the affine transformation implemented in region *r* lies from its local least-squares solution. Because the same matrices *W*_*l*_ are shared across regional transformations, these deviations are not independent, and their weighted contributions must cancel collectively. In this sense, a nonlinear network at stationarity globally balances regional errors, with Δ_*r*_ quantifying the effect of shared weights on regional least-squares optimality. In the single-region linear setting, there is no coupling between regions, and stationarity recovers the least-squares solution, Δ = 0 [1, 2]. We next investigate when the deviations Δ_*r*_ also become negligible in nonlinear networks.

#### Notations

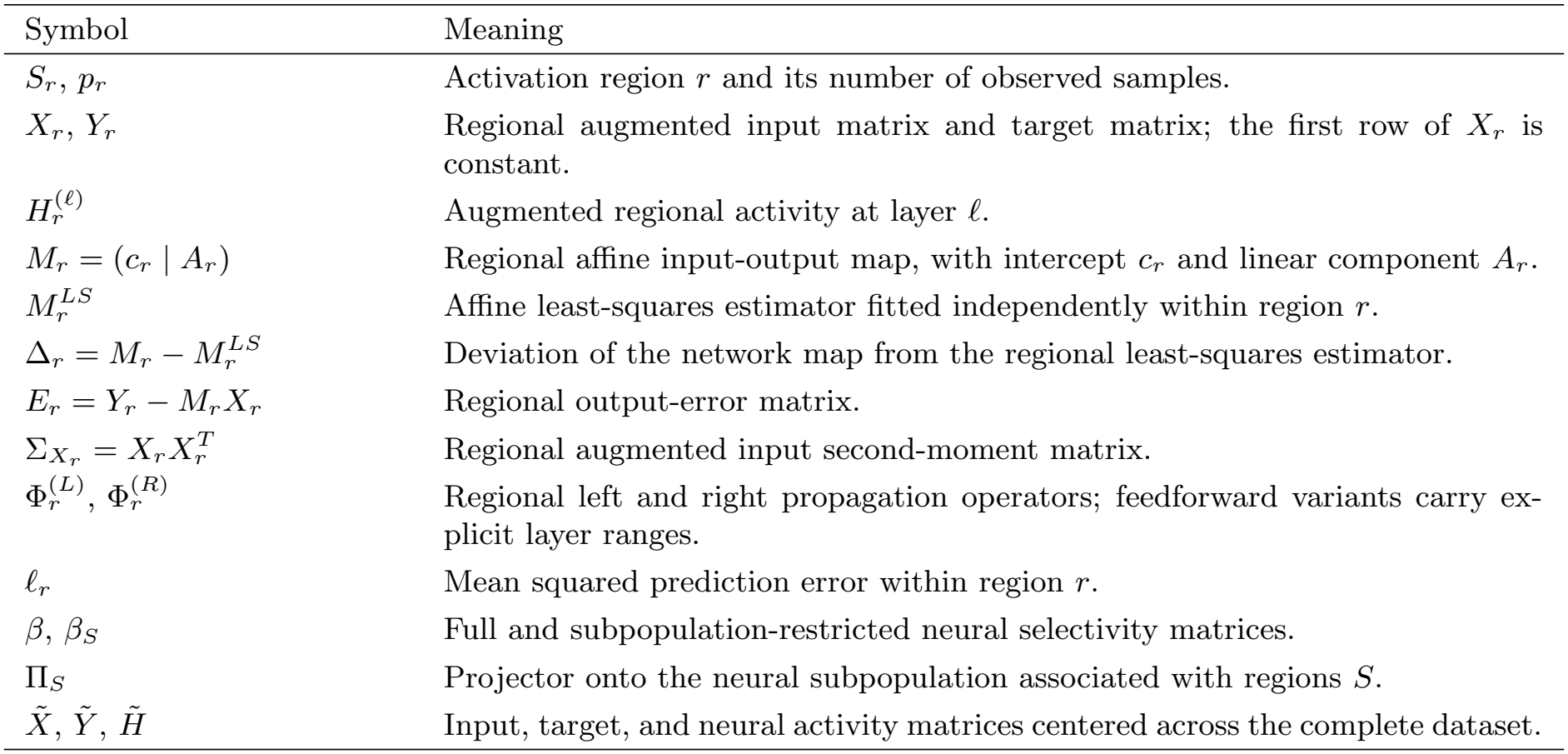

| Symbol | Meaning |
|---|---|
| S<sub>r</sub>, p<sub>r</sub> | Activation region r and its number of observed samples. |
| X<sub>r</sub>, Y<sub>r</sub> | Regional augmented input matrix and target matrix; the first row of X<sub>r</sub> is constant. |
| H<sub>r</sub><sup>(ℓ)</sup> | Augmented regional activity at layer ℓ. |
| M<sub>r</sub> = (c<sub>r</sub> \| A<sub>r</sub>) | Regional affine input-output map, with intercept c<sub>r</sub> and linear component A<sub>r</sub>. |
| M<sub>r</sub><sup>LS</sup> | Affine least-squares estimator fitted independently within region r. |
| Δ<sub>r</sub> = M<sub>r</sub> − M<sub>r</sub><sup>LS</sup> | Deviation of the network map from the regional least-squares estimator. |
| E<sub>r</sub> = Y<sub>r</sub> − M<sub>r</sub>X<sub>r</sub> | Regional output-error matrix. |
| Σ<sub>X<sub>r</sub></sub> = X<sub>r</sub>X<sub>r</sub><sup>T</sup> | Regional augmented input second-moment matrix. |
| Φ<sub>r</sub><sup>(L)</sup>, Φ<sub>r</sub><sup>(R)</sup> | Regional left and right propagation operators; feedforward variants carry ex-plicit layer ranges. |
| ℓ<sub>r</sub> | Mean squared prediction error within region r. |
| β, β<sub>S</sub> | Full and subpopulation-restricted neural selectivity matrices. |
| Π<sub>S</sub> | Projector onto the neural subpopulation associated with regions S. |
| X̃, Ỹ, H̃ | Input, target, and neural activity matrices centered across the complete dataset. |

## 3 Convergence toward regional least-squares solutions

We now consider expert networks that achieve low prediction error within each activation region. As established above, shared network parameters can shift each regional affine transformation away from its local least-squares solution. We now demonstrate that this deviation vanishes as the regional prediction error decreases, so that each regional predictor converges to its corresponding least-squares solution. Assuming, as above, that *X*_*r*_ has full row rank, define the error as

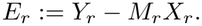

Then

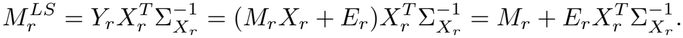

Hence

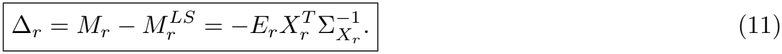

Let *p*_*r*_ = |*S*_*r*_| denote the number of samples in region *r*. Let 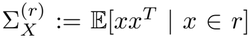 denote the population second-moment matrix of the augmented input distribution conditional on region *r*. This matrix captures the regional input statistics independently of any particular finite sample realization. For i.i.d. augmented inputs within region *r*, the law of large numbers gives

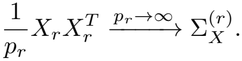

Since 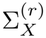 is symmetric positive definite, define its smallest eigenvalue as 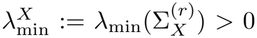. We then obtain

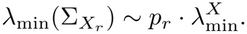

Because 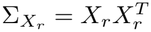, the singular-value decomposition of *X*_*r*_ gives

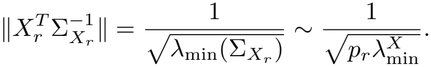

Where ∥ · ∥ denotes the spectral norm, Eq. (11) therefore gives

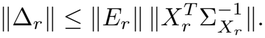

We bound the spectral norm of *E*_*r*_ by its Frobenius norm and define the average per-sample loss in region *r* as

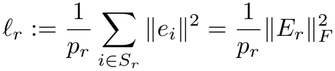

so that

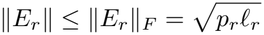

Combining these bounds gives

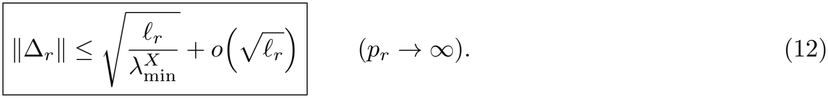

The dependence on 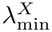 expresses the role of regional input conditioning: the same prediction error permits a larger deviation from least squares when the samples poorly constrain some input directions inside a region. By Eq. (12), the deviation from the affine least-squares solution vanishes whenever the per-region training loss vanishes.

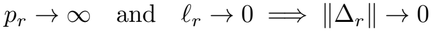

with rate 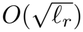.

In other words, low training loss within a region forces the network to behave like the optimal affine transformation on that region’s data. This result requires variation along every input direction within the region. If *X*_*r*_ is rank deficient, convergence can be established only on the subspace spanned by the regional inputs.

## 4 Tree interpretation of nonlinear networks and empirical validation of results

The previous results imply that piecewise-linear networks admit a simple functional interpretation. Since the network implements an affine transformation within each activation region, its global computation can be functionally decomposed into two stages: first, the input is routed toward an activation region through a hierarchical partition of the input space; second, a region-specific affine transformation is applied to produce the output. This interpretation is illustrated in Fig. 2a. Under this view, a piecewise-linear network is functionally equivalent to a decision tree whose leaves correspond to activation regions *r*, each associated with a local affine transformation *M*_*r*_. Related interpretations of piecewise-linear feedforward networks as decision trees have been described previously [17, 18]. Our analysis further shows that learning drives these regional transformations toward the corresponding affine least-squares solutions,

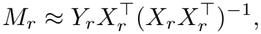

so that each leaf of the tree behaves as a locally optimal affine transformation specialized to the input-output statistics of its region.

**Fig. 1.**
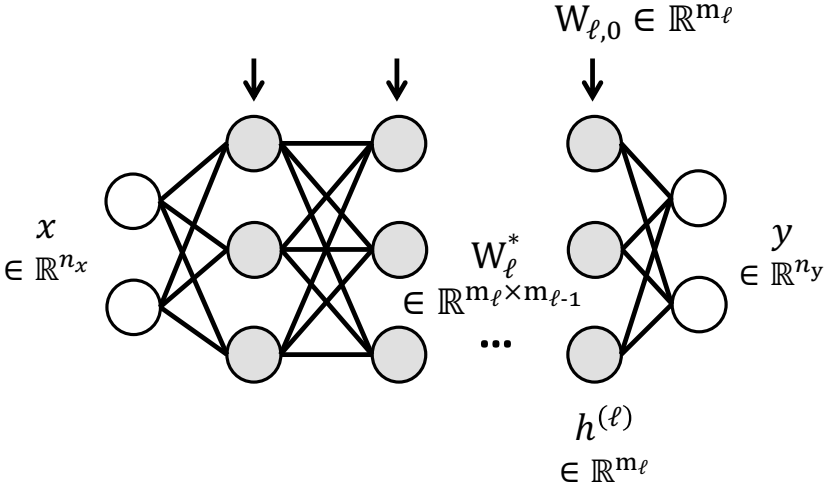
Schematic of the feedforward model analyzed in this section

**Fig. 2.**
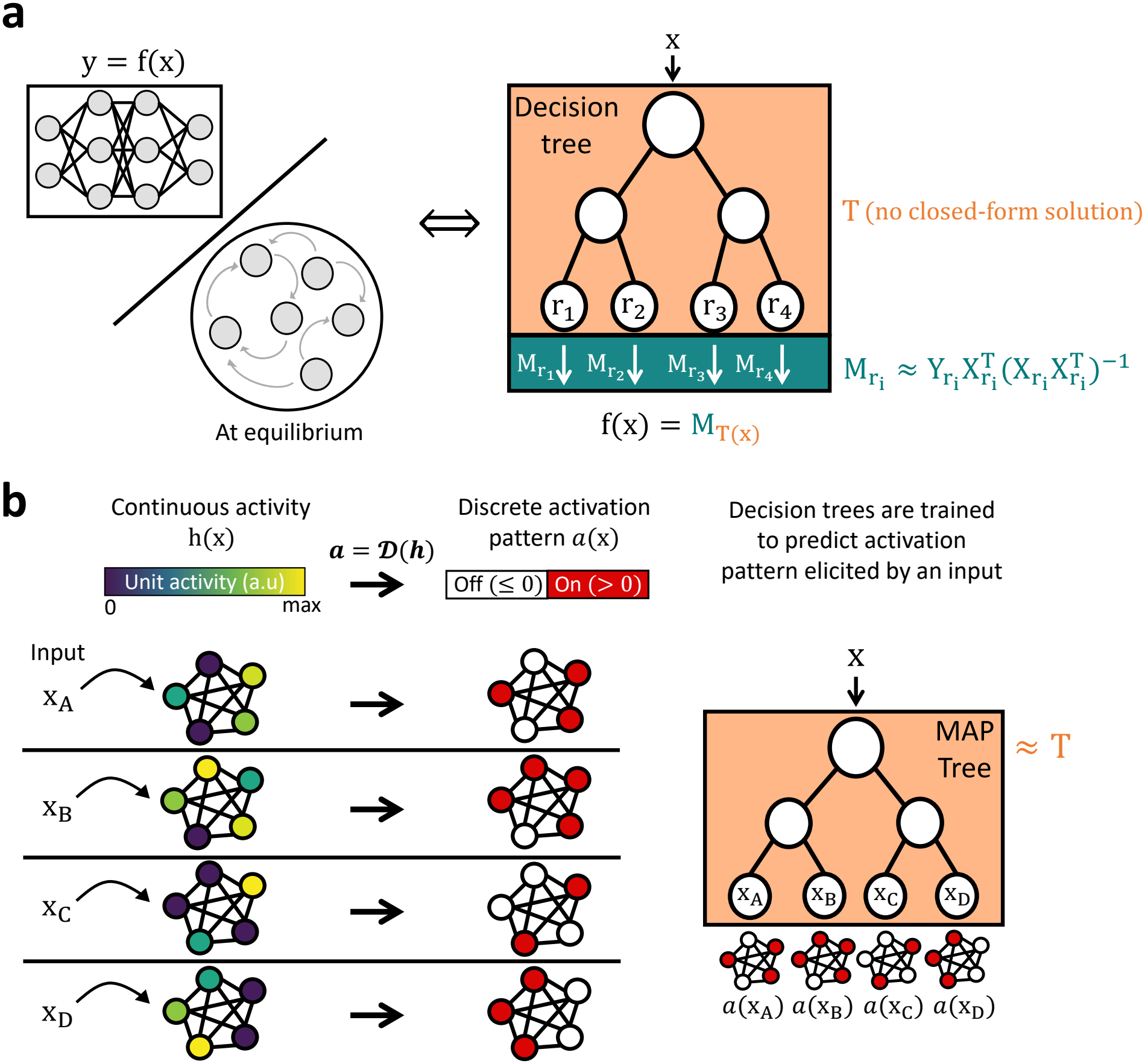
Tree representation of piecewise-linear neural computations. **a**, A piecewise-linear feedfor-ward network, or a recurrent network evaluated at equilibrium (see below), can be decomposed into a routing function *T* (*x*) that assigns each input *x* to an activation region *r*, followed by the corresponding regional affine transformation *M*_*r*_. Although the routing function has no general closed-form expression, the regional transformation approaches the local least-squares solution 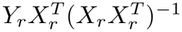 in the low-error regime. **b**, Numerical approximation of the routing function. A discretization function *D* converts continuous activity *h*(*x*) into a binary pattern *a*(*x*) indicating whether the activation function of each unit has a zero or nonzero local derivative. For the ReLU example shown here, these states correspond to inactive (*h* ≤ 0) and active (*h >* 0) units, respectively. A Main Activation Pattern (MAP) tree is then trained to predict the activation pattern *a*(*x*) from the input *x*, providing an explicit approximation of *T* (*x*) and assigning inputs to activation-defined regions.

Importantly, the derivations above characterize the computation performed *within* each activation region, but do not provide a closed-form description of how these regions are distributed across the input space. To make practical use of the theoretical predictions, the underlying routing process must therefore be approximated numerically. Because each activation region is defined by its activation pattern, a natural approximation of the latent routing structure is obtained by explicitly predicting these patterns from the input. We therefore trained decision trees to predict the dominant activation pattern elicited by a stimulus, yielding what we term *Main Activation Pattern* (MAP) trees (Fig. 2b). For artificial ReLU networks, units are active whenever their rectified activity is positive and inactive otherwise, so the binary vector directly identifies the corresponding activation region. For recorded neural activity, which has no known ReLU-like absolute threshold, activity states are instead defined relative to each neuron’s mean firing rate, as detailed in Methods. The MAP tree approximates the hidden partition structure implemented by the network by learning the mapping from inputs to activation patterns, thereby providing an explicit approximation of the latent decision process underlying nonlinear computation.

We first tested the regional least-squares prediction in feedforward networks trained to approximate a non-linear function of two continuous inputs (Fig. 3a). At successive stages of learning, we fitted MAP trees with increasing numbers of leaves and compared the affine map implemented in each inferred region with the corresponding local least-squares solution (Fig. 3b). This analysis tested both whether regional transformations approach least squares during learning and how finely this decomposition can be resolved by the network.

**Fig. 3.**
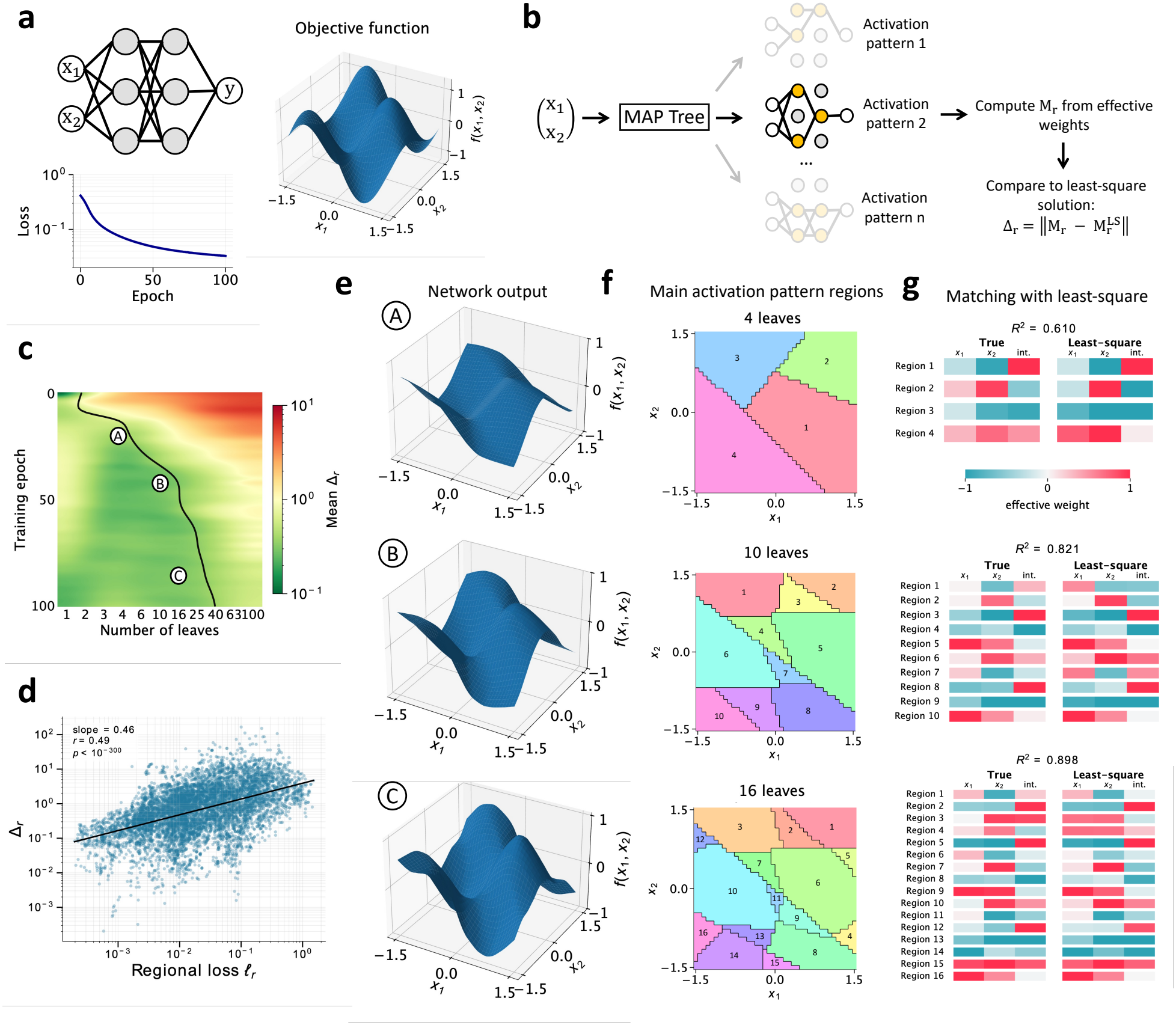
Convergence of regional transformations toward least-squares solutions in a feedforward network. **a**, Feedforward network trained to approximate a nonlinear two-dimensional target function; the training loss decreases over epochs. **b**, Analysis procedure. At each training stage, a MAP tree predicts the activation pattern from (*x*_1_, *x*_2_) and partitions the input space into regions. For each region *r*, the affine transformation *M*_*r*_ is computed from the network’s effective weights and compared with the regional least-squares solution 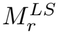 through the magnitude of the deviation, 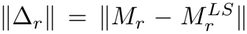. **c**, Mean regional deviation magnitude ∥Δ_*r*_∥ as a function of training epoch and number of MAP-tree leaves. The black trajectory marks the critical tree depth beyond which Δ_*r*_ increases, illustrating the finest granularity of the least-squares solutions captured by the network during learning. A-C indicate the stages illustrated in **e-g. d**, Regional deviation magnitude plotted against regional loss across training stages and MAP-tree regions. The log-log relationship has slope 0.46 (*r* = 0.49, *p <* 10^−300^), close to the square-root dependence predicted by the bound on ∥Δ_*r*_∥. **e**, Network output surfaces at stages A-C. **f**, Corresponding MAP-tree partitions of the input space into 4, 10, and 16 leaves. Colors identify distinct regions, and numbers denote the region indices used in **g. g**, Regional coefficients of the transformations implemented by the network (True; columns *x*_1_, *x*_2_, and intercept) and their least-squares counterparts for the partitions in **f**. Agreement increases as training progresses and regional error decreases (*R*^2^ = 0.610, 0.821, and 0.898 for A-C, respectively).

**Fig. 4.**
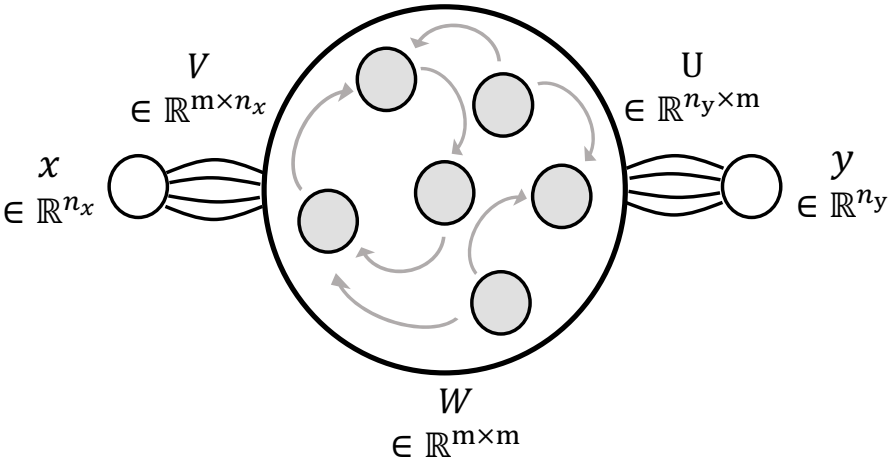
Schematic of the recurrent model analyzed in this section

Regional deviations decreased over training for a broad range of tree sizes (Fig. 3c). For each training stage, however, increasing tree depth beyond a critical value caused the deviation to rise again, identifying the finest partition at which the learned regional maps remained well described by local least squares. Across training stages and regions, the deviation scaled with regional loss with a log-log slope of 0.46, close to the square-root dependence predicted analytically (Fig. 3d). The same progression was visible directly in the learned surfaces and regional coefficients: agreement between network and least-squares transformations increased from *R*^2^ = 0.610 to 0.898 across the three illustrated stages (Fig. 3e-g). Thus, learning progressively resolved the nonlinear target into finer regional regressions.

## 5 Recurrent neural network at equilibrium

We next extend the regional least-squares framework to recurrent neural networks at equilibrium. A recurrent network admits a fixed-point representation that, upon unfolding, is formally equivalent to a deep feedforward network with tied affine weights. This equivalence allows us to carry the least-squares characterization over to the recurrent setting. The recurrent setting is of particular interest because it provides a natural model of cortical dynamics, where neurons interact through reciprocal connections and settle into a steady state in response to a sustained input. The aim of this section is therefore to yield predictions that are directly testable against electrophysiological recordings in biological neural networks.

### 5.1 Setup

For a sustained ordinary input 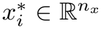, the network state *κ*_*i*_(*t*) ∈ ℝ^*m*^ evolves according to the continuous-time dynamics

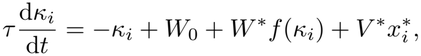

where *τ* is the neural time constant, *W*_0_ is the recurrent bias, *W* ^*^ is the recurrent weight matrix, *V* ^*^ is the input projection, and *f* is a coordinate-wise piecewise-affine activation function. We focus on stable equilibria of these dynamics, obtained by setting d*κ*_*i*_*/*d*t* = 0, and henceforth let *κ*_*i*_ denote the equilibrium state elicited by input 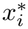. As in the feedforward network, the piecewise-affine activation partitions the inputs according to the affine piece selected at equilibrium. Writing *f*_*i*_ for the piece selected by input *i*, the recurrent activation regions are

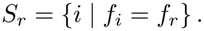

Within region *r*, the activation has a fixed diagonal slope matrix *D*_*r*_ ∈ ℝ^*m*×*m*^ and a fixed intercept.

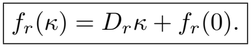

For ReLU, *D*_*r*_ is a diagonal binary mask and *f*_*r*_(0) = 0.

We retain the augmented-coordinate convention introduced above and write

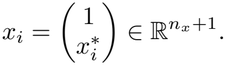

Thus,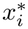denotes the ordinary input, whereas *x*_*i*_ denotes its augmented representation. The recurrent and input weight matrices are

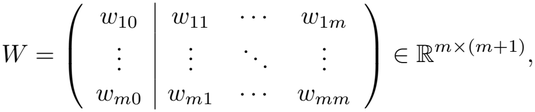

and

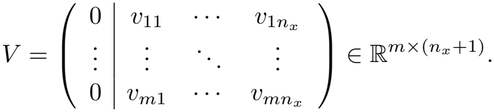

In compact block notation,

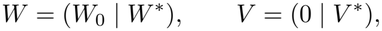

where *W*_0_ is the trainable recurrent bias, *W* ^*^ is the recurrent weight matrix, and *V* ^*^ is the input weight matrix. The first column of *V* vanishes because the constant input coordinate is already represented in *W*. Defining

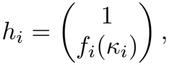

the equilibrium equation for the full recurrent network is

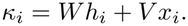

For *i* ∈ *S*_*r*_, this becomes

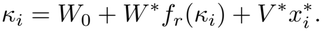

Let *b*_*r*_ denote the regional preactivation offset. It satisfies

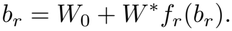

Define

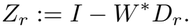

Subtracting the regional offset equation from the equilibrium equation gives

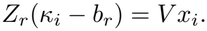

If *Z*_*r*_ is invertible, the regional fixed point is therefore

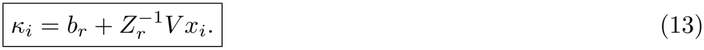

The inverse 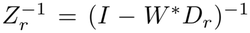 is the recurrent counterpart of a feedforward propagation operator. It captures the cumulative effect of recurrent interactions on an input perturbation within region *r*.

### 5.2 Gradient and stationarity condition

For the readout *ŷ*_*i*_ = *Uf*_*r*_(*κ*_*i*_), with 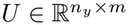, the regional map is affine. Since

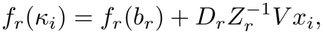

defining

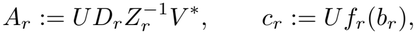

the regional readout takes the affine form

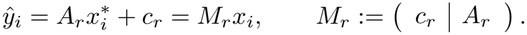

Differentiating the fixed-point equation gives 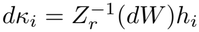 and hence

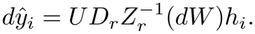

Defining the sample error *e* = *y* − *M x* and the left propagation operator 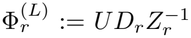, the gradient becomes

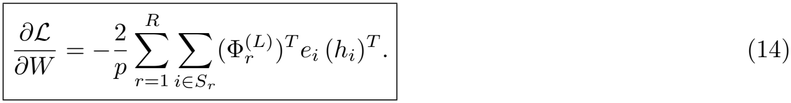

To express the activity term *h*_*i*_ in terms of the input, we similarly define the augmented right propagation operator

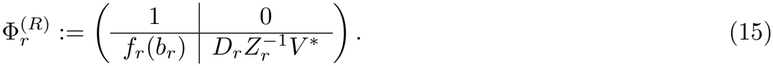

The equilibrium activity therefore satisfies

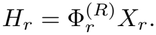

Setting *∂ ℒ /∂W* = 0 and using the regional least-squares relation in Eq. (9) gives the recurrent stationarity condition

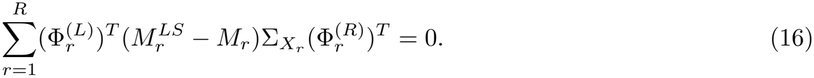

Thus

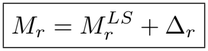

with deviation Δ_*r*_ meeting the constraint obtained from Eq. (16),

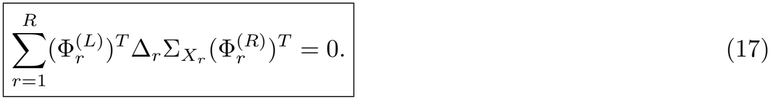

This condition has the same structure as its feedforward counterpart. The left operator propagates output errors through the recurrent dynamics, whereas the right operator maps inputs to equilibrium activities. Global stationarity therefore constrains a weighted sum of regional affine deviations, with the weights now determined by the recurrent interactions rather than a finite product of layer matrices.

#### Special case: fixed zero bias

If *W*_0_ = 0 and the activation maps zero to zero, then *b*_*r*_ = 0 and *c*_*r*_ = 0. The augmented stationarity condition then reduces to its non-augmented form.

Because the recurrent map has the same regional least-squares decomposition, the low-error convergence argument derived above applies without modification. We tested this extension in recurrent networks trained on a rule-switch task in which a contextual cue made either *x*_1_ or *x*_2_ relevant for the Go/NoGo decision (Fig. 5a). MAP trees fitted to equilibrium activation patterns recovered task-dependent partitions of stimulus space whose resolution increased with the number of leaves (Fig. 5b). We then asked whether these partitions were required to predict the equilibrium reached by the network. A single global linear fixed-point approximation systematically displaced the predicted equilibria from the true network states (Fig. 5c). In contrast, using the activation mask associated with each MAP-tree region brought the analytical fixed points close to the observed equilibria (Fig. 5d). This improvement held at the level of individual units: regional predictions closely followed the identity line in the examples shown, whereas the global approximation failed to capture their input-dependent activity (Fig. 5e). Across the population, incorporating the regional partition strongly reduced prediction error (Fig. 5f, *W* = 0, *p* = 1.82 × 10^−12^). These results establish that the regional decomposition extends from feedforward transformations to the equilibrium activity of recurrent networks.

**Fig. 5.**
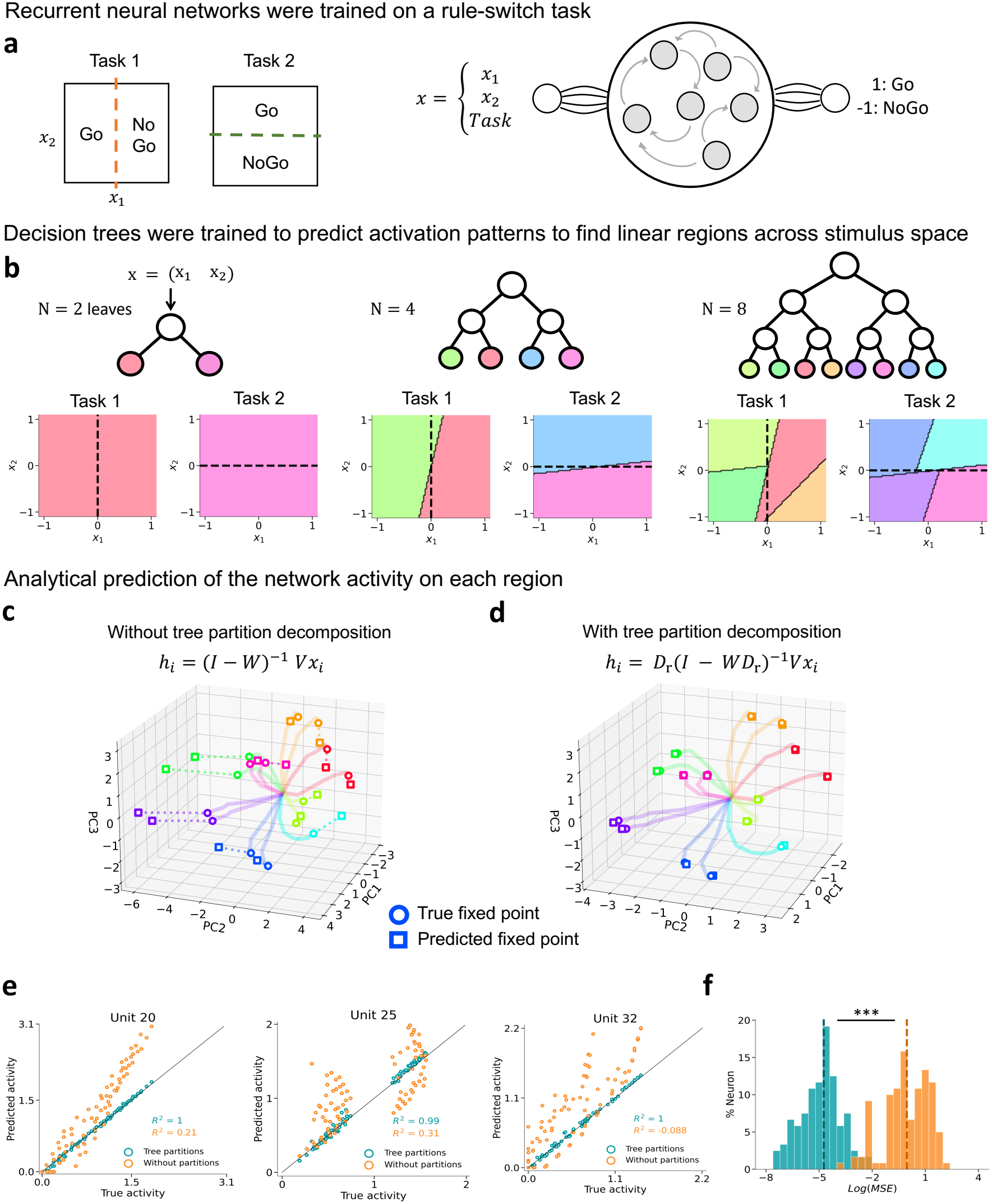
Activation-region decomposition predicts equilibrium activity in recurrent neural networks. **a**, Recurrent networks were trained on a rule-switch task in which the relevant decision boundary depends on a task cue: Task 1 separates Go and NoGo responses according to *x*_1_, whereas Task 2 separates them according to *x*_2_. The network receives (*x*_1_, *x*_2_, Task) and produces readouts for the binary Go/NoGo decision after reaching equilibrium. **b**, MAP trees with 2, 4, or 8 leaves were fitted to the equilibrium activation patterns. Colored areas show the resulting activation-defined partition of stimulus space separately for the two tasks; dashed lines show the task decision boundaries. **c**, Equilibrium activity predicted using a single global linear fixed-point approximation, without decomposing the input space into MAP-tree regions. True and predicted fixed points are shown in principal-component space by circles and squares, respectively; colors identify activation regions and trajectories show the network dynamics. **d**, Same comparison using the region-specific fixed-point solution determined by the activation mask *D*_*r*_. Accounting for the regional partition brings the predicted fixed points close to the true equilibria. **e**, Predicted versus true equilibrium activity for three example units. Blue points use the MAP-tree partition and orange points use the global approximation; the diagonal indicates equality and the displayed *R*^2^ values quantify prediction accuracy. **f**, Distribution across neurons of the log mean-squared error between predicted and true equilibrium activity, with and without the MAP-tree partition. Dashed vertical lines indicate the median of each distribution. Errors were lower with the MAP-tree partition (two-sided paired Wilcoxon signed-rank test across 40 neurons, *W* = 0, *p* = 1.82 × 10^−12^).

## 6 How neural selectivities are shaped by learning

We now apply the low-error result derived above to the geometry of neural selectivity. We consider a network with ReLU activation functions. In the large-sample, low-error regime, Δ_*r*_ → 0 and each regional affine map approaches its local affine least-squares solution. In this section, *X* and *H* denote ordinary, non-augmented input and activity matrices.

### 6.1 Linear encoding model

To characterize the geometry of neural selectivity, we consider the linear encoding model

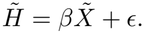

Here 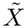 and 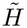 are centered across the complete dataset, and *β* contains the linear selectivity of every neuron to every input dimension. This is the standard encoding-model description used to quantify stimulus selectivity from neural recordings. Writing

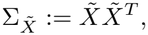

the least-squares estimator and its Gram matrix are

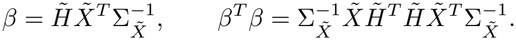

The matrix *β*^*T*^*β* describes the geometry of these selectivities in input space. Its diagonal entries measure the total selectivity associated with each input dimension, while its off-diagonal entries measure the alignment between selectivity directions. Characterizing this geometry therefore reduces to understanding the activity Gram matrix 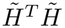.

### 6.2 Block decomposition over activation regions

Piecewise-linear networks partition the input space into activation regions. For the recurrent model, write

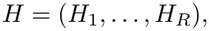

where 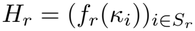 is the equilibrium activity in region *r*.

Define the global means

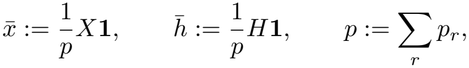

and the centered regional blocks

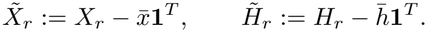

Writing

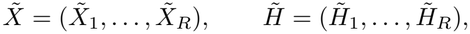

we obtain

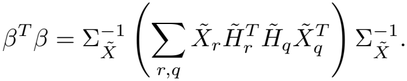

Thus the selectivity geometry decomposes into pairwise interactions between activation regions. Diagonal terms compare activity patterns within the same region, whereas off-diagonal terms measure how the representations associated with two different regions align.

### 6.3 Selectivity geometry after learning

For the targets, define

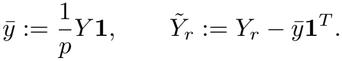

Within region *r*, the network implements the affine map

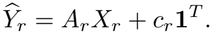

In the large-sample, low-error regime established above, this map approaches the corresponding affine least-squares solution. Its normal equations state that the residual has zero mean and is orthogonal to the inputs,

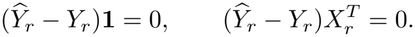

Since 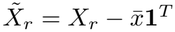, they imply

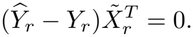

Summing (*Ŷ*_*r*_ − *Y*_*r*_)**1** = 0 over the regions gives *Ŷ* **1** = *Y* **1**, with *Ŷ* = (*Ŷ*_1_, …, *Ŷ*_*R*_) = *UH*, that is

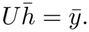

Therefore 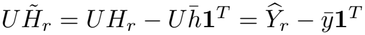, and

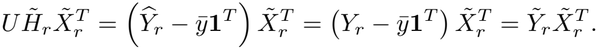

To relate this output-space constraint to neural activity, we assume that the readout matrix

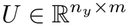

has normalized orthogonal rows, such that

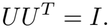

This relation does not uniquely determine 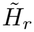 since activity components in ker(*U*) are invisible to the output. However, activity orthogonal to the readout subspace contributes no functional signal while still incurring metabolic cost. Under activity regularization, stationarity therefore favors minimum-energy representations concentrated in the row space of *U*. Under this assumption,

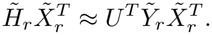

For two regions *r* and *q*, this representation gives

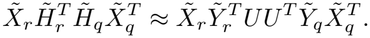

Using *UU* ^*T*^ = *I*, pairwise interactions between regional activity patterns are therefore determined by the alignment of their output-input covariance matrices. Defining

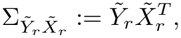

we obtain

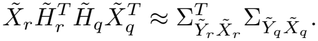

Inserting this into the block decomposition of *β*^*T*^ *β* above gives

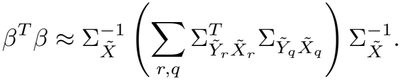

Defining

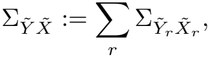

we obtain

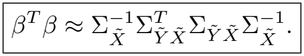

The global selectivity geometry is therefore inherited from the total output-input covariance. In this sense, a nonlinear network recovers the same covariance structure as a linear model after its region-specific solutions are aggregated.

### 6.4 Modular decomposition over neural subpopulations

We now characterize the geometry of selectivity over neural subpopulations. Let Π_*S*_ project onto neurons whose activation slope is nonzero only in regions belonging to *S*, and define *β*_*S*_ := Π_*S*_*β*. Within an activation region, each ReLU unit is either active, with nonzero slope, or inactive, with zero output. A neuron selected by Π_*S*_ is active only in regions belonging to *S* and is therefore inactive in every region *r* ∈*/ S*. Consequently,

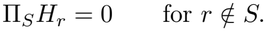

Since 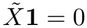, centering the activity does not change its covariance with the input.

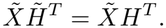

It follows that

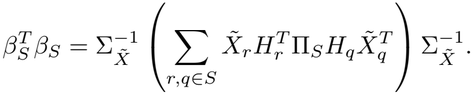

Thus neurons selected by Π_*S*_ contribute no activity outside the regions on which their activation slope is nonzero. The global sum over regions therefore reduces to interactions entirely within *S*. Using the minimum-energy representation, 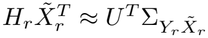 with 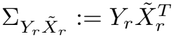, gives

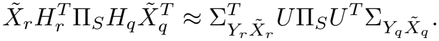

Assuming that the activation-defined subpopulation is sufficiently large and isotropic in readout space, so that

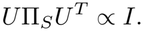

Then

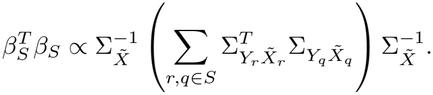

Defining

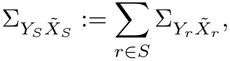

gives

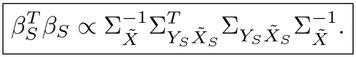

This assigns a distinct covariance geometry to each activation-defined neural subpopulation. Each subpopulation behaves as a local predictor specialized to the input-output statistics of the regions in *S*, while the outer factors 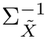 account for correlations in the complete stimulus distribution. Under input whitening, 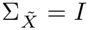, this reduces to 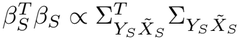.

### 6.5 Validation of selectivity predictions across simulated and empirical networks

We tested this prediction with three simulated recurrent networks performing distinct context-dependent tasks and two empirical datasets. Across the simulated tasks, context determined either which of two stimulus features was used for categorization (Supplementary Fig. 1), the mixture of features defining the category boundary (Supplementary Fig. 2), or whether the features were combined additively or multiplicatively (Supplementary Fig. 3). The empirical analyses used prefrontal recordings from mice switching between orientation and spatial-frequency discrimination [12] (Fig. 6) and neural recordings from mice using context to select between visual and auditory evidence [16] (Supplementary Fig. 4). Across all five analyses, MAP trees fitted to predict population activation patterns from the input recovered the task boundaries without access to target outputs (Supplementary Figs. 1c, 2c, 3c, and 4b; Fig. 6b). In the first empirical dataset, we analyzed 2,167 prefrontal neurons (Fig. 6a). Neurons were grouped according to the set of the regions in which they were active, producing nine subpopulations with distinct distributions in orientation, spatial-frequency and task-selectivity space.

**Fig. 6.**
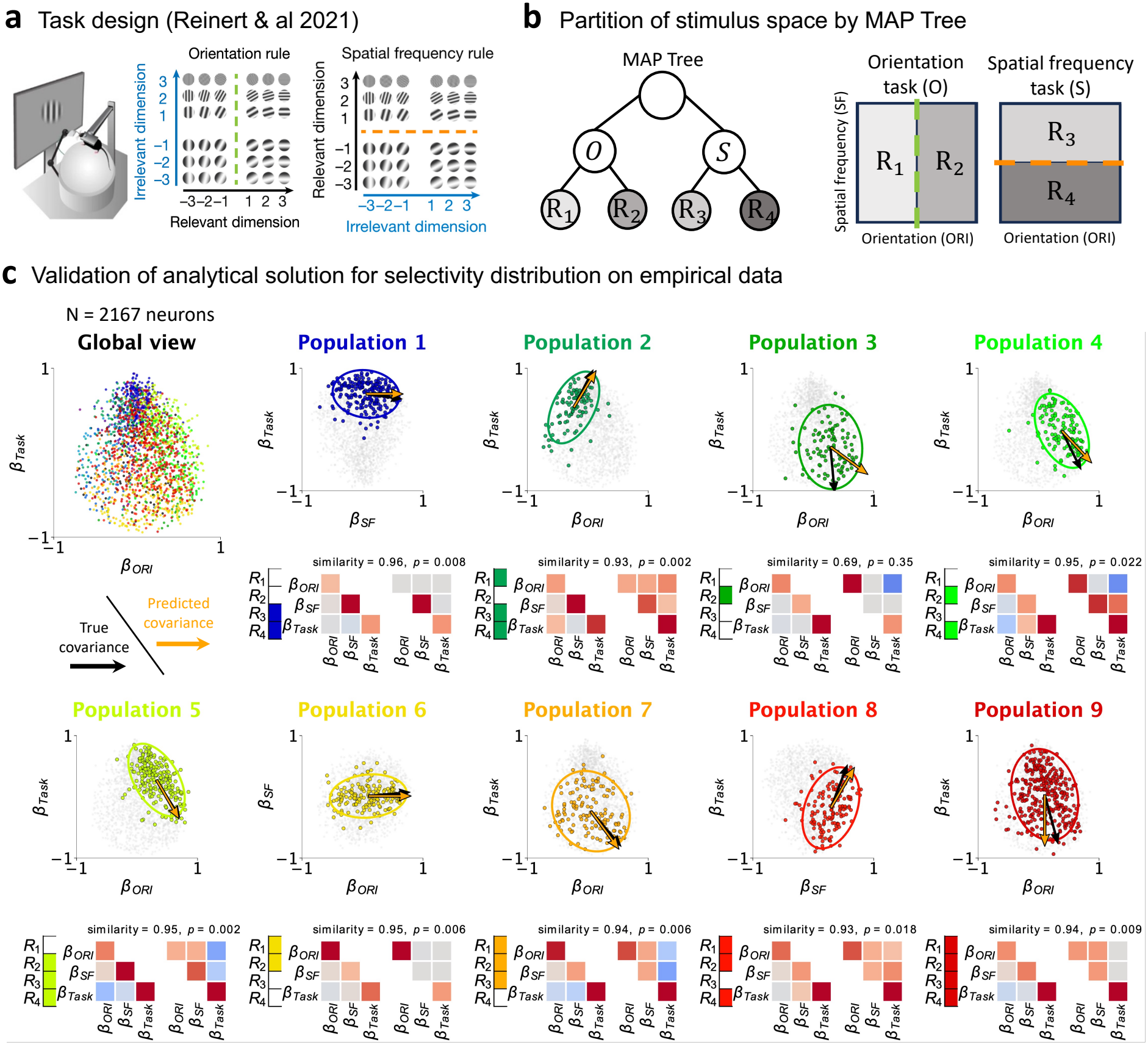
Task statistics predict the selectivity geometry of activation-defined prefrontal neural subpopulations. **a**, Task design adapted from Reinert et al. (2021). Mice discriminated stimuli according to either their orientation or their spatial frequency. Under the orientation rule, orientation was behaviorally relevant and spatial frequency was irrelevant; under the spatial-frequency rule, these roles were reversed. Green and orange dashed lines indicate the category boundary along the task-relevant stimulus dimension. **b**, Partition of the joint stimulus space by a MAP tree fitted to population activity. The first split separates the orientation (O) and spatial-frequency (S) task contexts; subsequent splits along the relevant stimulus dimension define four activation regions (*R*_1_, *R*_2_, *R*_3_, and *R*_4_), shown in the corresponding stimulus spaces. **c**, Validation of the predicted selectivity geometry in 2,167 recorded neurons. The global view shows neurons in selectivity space, and the nine detailed plots show activation-defined subpopulations corresponding to distinct combinations of MAP-tree regions. In each plot, colored points highlight the selected subpopulation against the remaining neurons in gray, and the ellipse represents its measured covariance in the displayed pair of selectivity dimensions. Black and orange arrows indicate the principal directions of the measured and theoretically predicted covariance, respectively. The adjacent *R*_1_-*R*_4_ indicators mark in color the activation regions defining each subpopulation. Covariance matrices compare the measured lower triangle with the theoretically predicted upper triangle across orientation, spatial-frequency, and task selectivities. Matrix entries are rescaled coefficients ranging from −1 (blue) to +1 (red). The displayed similarity is the cosine similarity between the unique entries of the measured and predicted covariance matrices; the associated *p* value is obtained from the rotation-based permutation test described in Methods.

For each subpopulation, we predicted selectivity geometry from the input-output statistics pooled over its associated regions and compared this prediction with the covariance measured from neural activity (Fig. 6c; Supplementary Figs. 1d, 2d, 3d, and 4c). Selectivity geometry varied markedly across subpopulations, but this variation was accurately captured by the task-derived predictions, which recovered both the covariance structure and its principal directions. The mean similarity across subpopulations was significant in all five analyses (*p <* 10^−3^ in each). Together, these results establish activation-region structure as the basis of modular selectivity. Neurons active across different sets of regions form distinct subpopulations whose geometry reflects the input-output statistics processed within those regions.

## 7 Neural baseline controls the balance between coarse- and fine-grained learning regimes

Our tree interpretation directly links single-neuron operating points to network-level expressivity. We use neural baseline to denote the input-independent bias that sets unit preactivations relative to the ReLU threshold. By determining how readily units cross this threshold as inputs vary, neural baseline controls the emergence of new activation patterns and, in turn, the number of distinct affine maps the network can implement.

When neural baselines lie far above threshold, most units remain active for all inputs (Fig. 7, left). Their slope matrices are then nearly constant across the input space, so the network visits only a small number of activation regions. In the limiting case in which every unit remains on the same linear branch, *D*_*r*_ = *I* and the regional decomposition collapses to a single affine map. The corresponding MAP tree has one leaf, and the network implements a single regression plane. Despite having many parameters, its effective nonlinear expressivity is therefore low.

**Fig. 7.**
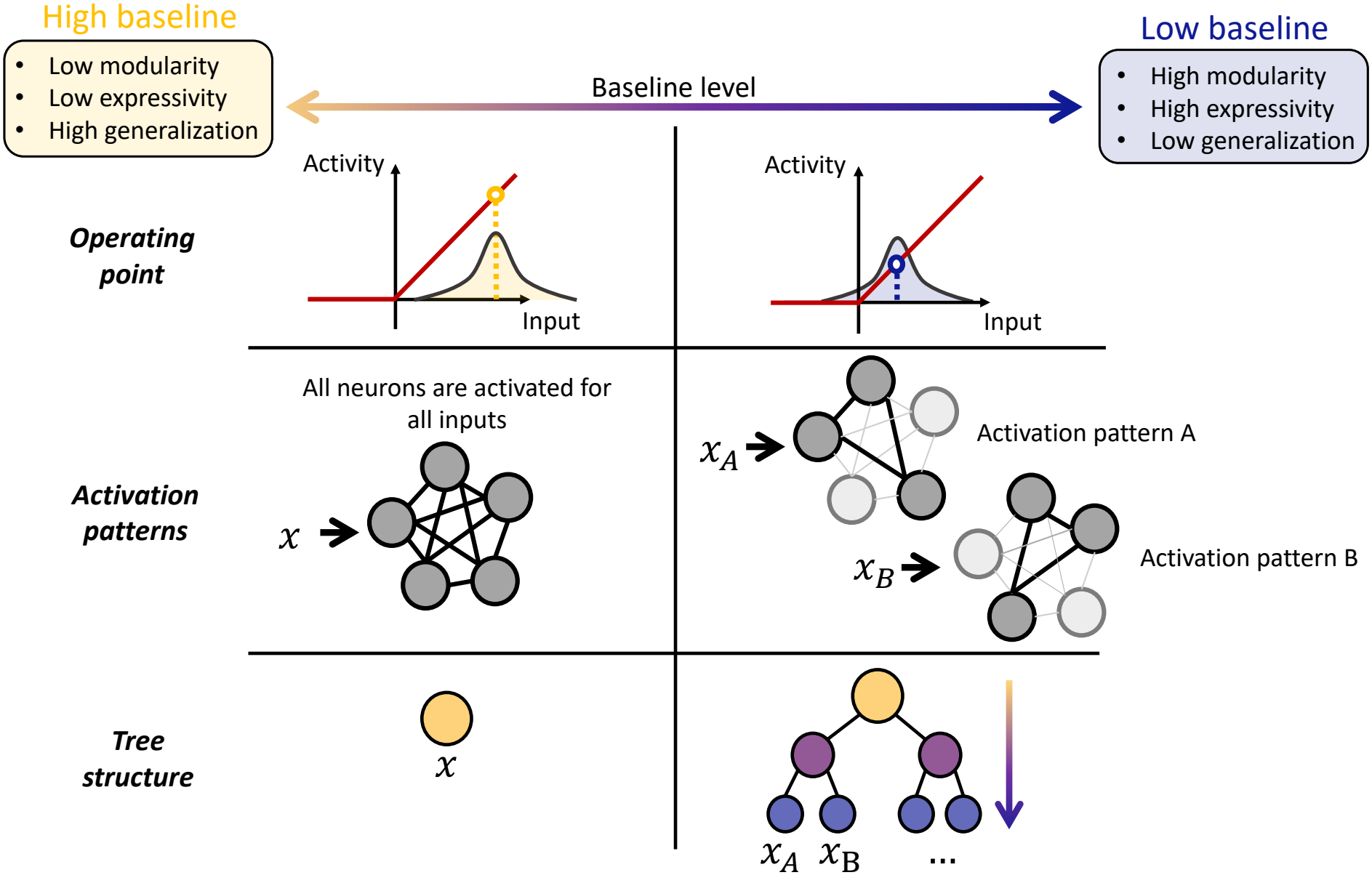
Neural baseline controls activation pattern diversity and the resolution of learned input-output structure. The horizontal continuum indicates a decrease from a high neural baseline (left, yellow) to a low baseline closer to the activation threshold (right, blue). At a high baseline, the distribution of stimulus-driven inputs lies primarily on the active branch of the rectified-linear activation function. The same units therefore remain active across inputs, producing a common population activation pattern and routing the inputs to a single tree region. This regime implements few regional transformations and is predicted to have low modularity and expressivity but stronger generalization. At a lower baseline, the stimulus distribution straddles the activation threshold, so changes in the input switch subsets of units between active and inactive states. Different inputs, illustrated by *x*_*A*_ and *x*_*B*_, consequently recruit distinct population activation patterns and are routed toward different leaves of a deeper tree. The downward arrow denotes the continued growth of the regional hierarchy as additional patterns emerge. This regime is predicted to support greater modularity and expressivity at the cost of reduced generalization. Red curves show the common rectified-linear activation function; dashed vertical lines and colored circles indicate the operating point in each regime; filled and unfilled neurons denote active and inactive units, respectively.

Moving the population baseline toward threshold has the opposite effect. Small changes in the input can now switch individual units between their inactive and active states, producing a larger collection of activation patterns across the input space (Fig. 7, right). These additional patterns appear as new leaves in the effective tree implemented by the network, each supporting a distinct local affine map.

This gain in expressivity for low-baseline networks may, however, come at a statistical cost. Every additional leaf partitions the training set more finely, leaving fewer examples to constrain each regional affine map. Finer regional partitions therefore support highly specialized solutions, but these solutions are estimated from smaller effective sample sizes and are more sensitive to the particular examples assigned to each region. New inputs may consequently be assigned to local models that received little nearby training data. In this finite-data regime, increasing activation-pattern diversity can improve performance on the training set while reducing the stability of predictions on unseen examples.

By contrast, a high-baseline network occupies fewer activation regions and pools more examples when estimating each local map. Its shallower internal tree has lower expressivity but imposes greater sharing across the dataset, favoring broad regularities over solutions specific to individual neighborhoods. This should reduce estimation variance and improve generalization when data are limited or when the target function is sufficiently smooth. The same constraint can nevertheless introduce approximation bias when the task requires genuinely different computations in different parts of the input space. Neural baseline should therefore control a bias-variance trade-off: moving toward threshold increases regional specialization, whereas moving away from threshold favors simpler solutions supported by more observations. The best operating point should depend jointly on task nonlinearity and the amount of available training data.

We test these anticipated effects of neural baseline on network expressivity and generalization below.

### 7.1 Activation-region formation enables nonlinear learning

We first examined how neural baseline affects network expressivity during learning. While a network remains confined to a single activation pattern, it implements one affine map and cannot improve beyond the global least-squares solution. Reducing the loss below this limit requires a new activation pattern and the additional regional transformation that it supports.

To test this prediction, we trained networks at different baselines and benchmarked their learning trajectories against two reference models: a one-plane solution fitted jointly across task contexts and a two-plane solution fitted separately within each context (Fig. 8, top). Learning unfolded in discrete stages that matched changes in activation-region structure (Fig. 8, bottom). At the highest baseline, the network occupied a single region and its loss plateaued near the one-plane solution. At an intermediate baseline, the emergence of a second occupied region coincided with the loss falling below the one-plane solution and approaching the two-plane solution. Finally, at the lowest baseline, the formation of a third region coincided with the loss falling below the two-plane limit. These trajectories show that the emergence of new activation regions enables successive reductions in loss beyond the limits of simpler affine solutions.

**Fig. 8.**
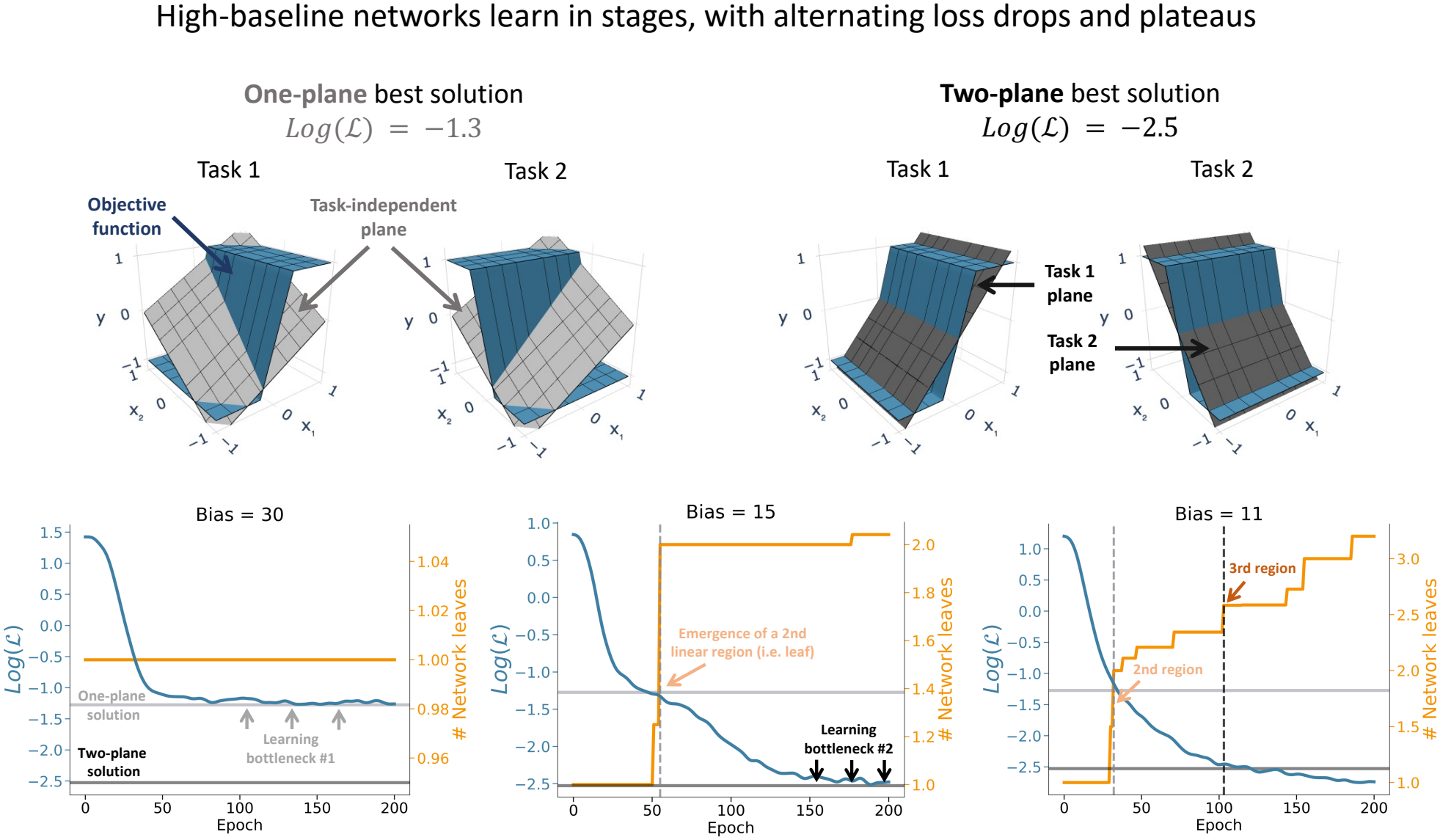
The emergence of activation regions enables learning beyond the global affine solution. **Top**, Geometric reference solutions for the rule-switch task. Blue surfaces show the task-dependent target transformation and gray surfaces show the best affine approximation. A single task-independent plane fitted jointly across both contexts yields the one-plane solution, with log(ℒ) = −1.3. Fitting one plane per task context yields the two-plane solution, with log(ℒ) = −2.5. **Bottom**, Learning trajectories for representative recurrent networks initialized with high (*b* = 30), intermediate (*b* = 15), or lower (*b* = 11) baselines. Blue curves show the logarithm of the training loss (left axes), and orange step functions show the mean number of occupied MAP-tree leaves across epochs (right axes). Gray horizontal lines indicate the errors of the one- and two-plane reference solutions. At *b* = 30, the network remains within one activation region and its loss plateaus near the one-plane solution. At *b* = 15, the appearance of a second activation region (dashed line) permits the loss to cross the one-plane boundary and approach the two-plane solution. At *b* = 11, successive appearances of second and third regions (dashed lines) are followed by additional decreases in loss. Gray and black arrows mark learning plateaus or bottlenecks. Together, the examples show that improvements beyond an affine reference solution coincide with increases in the number of regional transformations available to the network.

### 7.2 Fine-grained learning comes at a generalization cost

To assess how neural baseline affects generalization, we trained networks with different baselines to approximate the same nonlinear radial function from a sparse set of inputs distributed across concentric circles (Fig. 9a). Generalization was evaluated on a dense grid of previously unseen inputs spanning the full space. Networks operating closer to threshold fitted the training samples more accurately, whereas high-baseline networks achieved lower loss on the dense test grid (Fig. 9b,c).

**Fig. 9.**
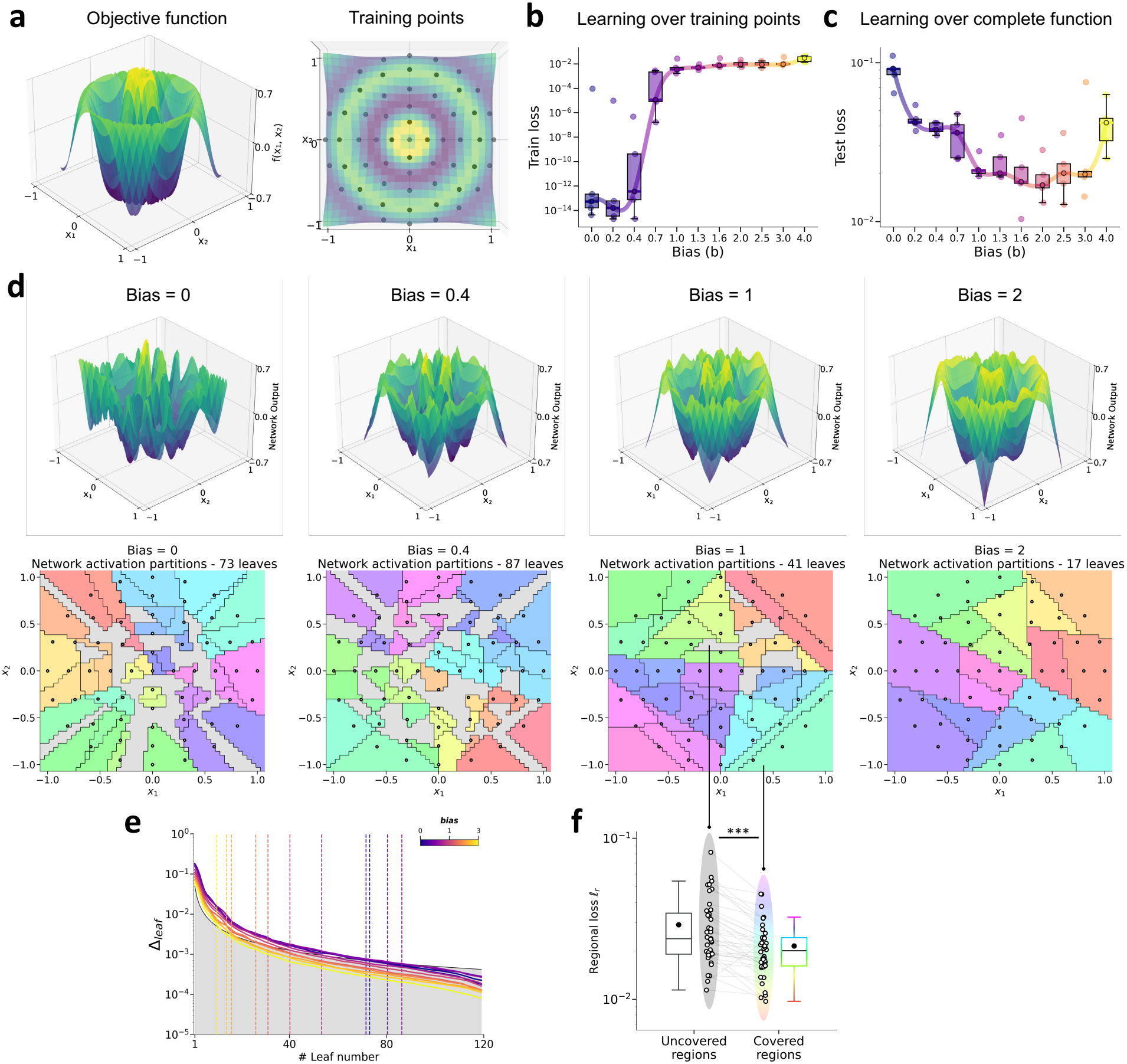
Neural baseline controls a trade-off between training fit and generalization. **a**, Nonlinear radial target function (left) and finite training set (right). Black points indicate the training inputs distributed over six concentric circles; the colored surface shows the target value. **b**, Final training loss as a function of neural baseline. The colored curve interpolates the median across baselines. Colors progress from low (purple) to high (yellow) baseline. Low-baseline networks achieve the smallest training losses. **c**, Final test loss on a dense held-out grid. In contrast to the training loss, high-baseline networks generalize better to these new samples. **d**, Final network output surfaces (top) and corresponding MAP-tree partitions of input space (bottom) for baselines *b* = 0, 0.4, 1, and 2. Colors in the partition plots identify distinct leaves, black points show the training inputs, and the number of occupied leaves is reported above each partition. Networks operating closer to threshold form finer partitions and reproduce smaller-scale features of the target, whereas higher-baseline networks form coarser partitions and smoother output surfaces. **e**, Marginal decrease in MAP-tree cross-entropy, Δ_leaf_, produced by each additional leaf. Curves show the mean across five networks for each baseline; colored dashed lines indicate the estimated effective number of leaves, defined by the point at which the incremental improvement falls below the depth-dependent threshold (black line; gray area). **f**, Regional loss, *l*_*r*_, within MAP-tree regions containing no training input (uncovered regions) and regions containing at least one training input (covered regions). Mean losses are larger in uncovered than covered regions (two-sided paired Wilcoxon signed-rank test, *n* = 55 networks, *W* = 29, *p* = 3.16 × 10^−9^), demonstrating the generalization cost of finely partitioning a finite training set. In **b, c**, and **f**, points denote individual network fits, boxes show the interquartile range and median, and black circles show the mean.

The learned functions and their MAP-tree partitions clarified why baseline had opposite effects on training and test performance (Fig. 9d). Low-baseline networks divided the input space into many regions and captured small-scale variations of the target, whereas increasing baseline reduced the effective number of leaves and produced smoother approximations. Consistent with this interpretation, MAP trees fitted to low-baseline networks required more leaves to capture activation patterns than those fitted to high-baseline networks (Fig. 9e). The finer partitions formed by low-baseline networks created regions containing no training samples, which we refer to as uncovered regions. Prediction error was significantly higher in uncovered than covered regions (*W* = 29, *p* = 3.16 × 10^−9^; Fig. 9f), indicating that poor generalization was concentrated in regions not visited by any training input. Together, these results show that neural baseline sets the resolution of learning: lower baselines favor fine-grained representations, whereas higher baselines favor generalization beyond the training data.

## 8 Discussion

### Inferring computation from representational geometry

A central contribution of our framework is to establish an analytical link between the function learned by a network and the observable geometry of its internal representations. Starting from the network objective, we derived necessary conditions on neural selectivity at learning stationarity. These constraints link each neuron’s selectivity to the input-output statistics of the regions in which it is active. Neurons active across different sets of regions therefore develop distinct mixtures of feature selectivity, giving rise to discrete subpopulations in selectivity space [9, 10, 13, 19]. Neural selectivity thus provides a measurable signature of the input-output relationships learned by the network. Consistent with this prediction, the selectivity geometry derived from task statistics closely matched the geometry observed across neural subpopulations in both simulated and empirical datasets.

Our analysis therefore establishes a forward mapping from task structure to neural representation. More broadly, the potential of this framework lies in *reversing* this mapping: observable properties of neural activity could be used to infer the underlying computations performed by a network. Given the modular organization of a population and the selectivity geometry of its constituent subpopulations, one could seek to recover the latent input-output relationships consistent with this geometry. This inverse problem will generally be underdetermined, as distinct input-output relationships can produce equivalent selectivity covariances, for example through rescaling or the influence of unobserved task variables. Nevertheless, the modular organization of selectivity carries substantial information about the underlying computation and sharply constrains the space of compatible functions. Extending the theory toward a full inverse formulation could therefore turn representational geometry into a tool for functional identification, providing a principled route from neural activity to the computations it implements.

### The dual structure of nonlinear computation

At a more abstract level, our results place nonlinear neural networks at the interface between discrete and continuous mathematics. Their activation structure partitions continuous input space into discrete regions, while an affine map specifies the input-output transformation within each region. These two operations are fundamentally different [4, 20]. The discrete component selects the rule, whereas the continuous component implements that rule. This distinction naturally separates input dimensions into modulators and drivers. Modulators, such as task or contextual cues, move the network across regional boundaries and switch the transformation being applied. Drivers, which may include stimulus features within a given region, continuously affect neural activity without changing the selected transformation [10, 14, 21]. In this sense, piecewise-linear networks derive their expressivity from operating at the boundary between combinatorial and continuous systems.

### Modularity in sensorimotor remapping tasks

We previously showed that sensorimotor remapping increases modularity in selectivity space [15]. The present framework provides a mechanistic explanation for this observation. In these tasks, the same sensory input must be mapped to different outputs depending on context, making the input-output relationship intrinsically nonlinear. To implement this nonlinear mapping, the network must switch between distinct activation regions across contexts. As a result, neurons recruited across different sets of activation regions become sensitive to different task statistics, and form distinct subpopulations in selectivity space. Under this view, modular selectivity emerges directly from the regional decomposition required to implement a nonlinear mapping. The resulting modular organization should therefore depend on how readily distinct activation regions form during learning.

### Neural baseline sets the resolution of learning

Our framework identifies neural baseline as a regulator of this process. By controlling activation-region diversity, baseline shapes the expressivity and modularity of learned representations and places networks along a continuum between two learning regimes. At high baseline, most units remain active across inputs, limiting the diversity of activation patterns and favoring a coarse representation of the input-output relationship. As baseline approaches the activation threshold, inputs recruit increasingly distinct neural populations, enabling a more modular and fine-grained representation. Our simulations support this transition from a “big-picture” learning regime, which captures broad task structure, to a “detail-oriented” regime capable of dealing with finer variations.

An important next step will be to test this prediction empirically. The theory suggests that global inhibitory or excitatory signals could shift biological networks between these learning regimes by modulating their operating baseline [22]. Tasks requiring fine discrimination of the input-output relationship should benefit from moderate inhibition that brings neural activity closer to threshold and increases the diversity of activation patterns. By contrast, tasks dominated by coarse, large-scale structure may benefit from a higher-baseline regime that limits unnecessary partitioning. These predictions directly link global modulation and representational modularity to the granularity of the input-output relationship that must be learned. Testing this prediction in biological networks will require a robust approach to identify activation regions from recorded activity.

### Extending activation-region analysis to biological dynamics

In the analytical framework, activation regions have an exact definition: they comprise inputs processed by the same local affine transformation. To translate this decomposition into a practical analysis, we focused on ReLU networks, for which each unit has only two derivative states. A unit is active when its preactivation is positive, with a nonzero derivative, and inactive otherwise, with zero derivative. MAP trees recover this local derivative structure by predicting the binary activation state of every unit from the input, thereby providing an explicit numerical approximation of the network’s activation regions.

Applying the same construction to biological data requires an additional assumption because the activation function of each recorded neuron is generally unknown. Neural activity must therefore be discretized into putative active and inactive states, interpreted as regimes with nonzero and zero local derivatives, respectively. This choice directly determines the inferred regional partition and its associated neural subpopulations. Establishing biologically grounded discretization rules, adapted to the response properties of the recorded cell types and circuits, will therefore be essential for strengthening the application of the framework to empirical neural data.

Defining these states does not, however, ensure that regional membership is stable across trials. Our derivation assumes deterministic, noise-free dynamics, whereby each input is assigned to a unique activation pattern and a corresponding region of input space. In biological networks, trial-to-trial variability breaks this one-to-one correspondence: repeated presentations of the same input can give rise to different population activity patterns and, consequently, different regional assignments. Activation regions should therefore be treated as stochastic rather than deterministic objects, characterized by input-dependent probabilities over activation patterns [23, 24]. Incorporating this variability into the framework, for instance through probabilistic region membership, will be an important step toward a more faithful description of biological neural computation.

Our recurrent-network derivation further assumes that neural activity reaches equilibrium for each input. This assumption may not hold in tasks with rapid temporal fluctuations, for which network activity continuously evolves without approaching a fixed point [25, 26]. The framework nevertheless extends naturally to slowly varying inputs. When the characteristic timescale of input variation is long relative to the network’s relaxation time, activity remains close to the instantaneous equilibrium associated with the current input. A time-dependent input-output relationship can then be treated as a sequence of quasi-stationary mappings, allowing the regional decomposition to be applied locally in time. Extending the theory to regimes in which input and network timescales are comparable will require an explicitly dynamical formulation. These extensions would describe variability around a regional partition, but leave open how that partition itself emerges and changes during learning.

### Dynamics of activation regions during learning

Our derivations characterize the affine transformation associated with each activation region while treating the regional partition as given. During learning, however, the transformations and the regions evolve together: changes in network parameters move regional boundaries and can create, merge, or eliminate regions. An analytical description of these dynamics would complete the present account of learning in nonlinear networks. In particular, it would reveal how initial weights and neural baselines determine the trajectories along which input space is partitioned, and why networks trained on the same task can converge to different regional decompositions despite achieving similar performance [12, 27].

Such a theory could also clarify variability in learning across individuals and training histories. Different initial conditions may bias learners toward distinct partitions of the same task, producing alternative strategies and representational geometries. The evolution of regions should also depend on how experience is ordered. During sequential learning, partitions established by earlier tasks provide a pre-existing structure that subsequent learning must reuse, refine, or reorganize, thereby shaping transfer and interference between tasks [11, 28]. Under joint or continuously interleaved learning, by contrast, regional boundaries can co-evolve with the statistics of all tasks. Describing these path-dependent regional dynamics would connect learning history to the diversity and constraints of the representations that ultimately emerge.

## 9 Methods

### 9.1 Network models and optimization

Networks used the rectified-linear activation *ϕ*(*z*) = max(0, *z*). The feedforward network comprised two hidden layers and was defined by

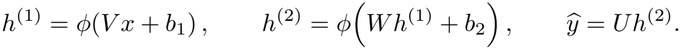

The recurrent networks received a constant input over the duration of each trial and followed the leaky discrete-time dynamics

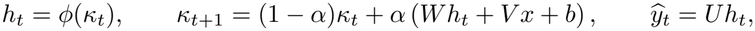

where *κ*_*t*_ is the recurrent state, *h*_*t*_ the rectified activity, *W* the recurrent matrix, *V* the input projection, *U* the readout, and *b* the additive recurrent bias. Because *b* determines the activity level of a unit in the absence of stimulus-driven fluctuations, we also refer to it as the neural baseline. We used *α* = 0.2. Activity and output were summarized over the terminal window of length *τ* as

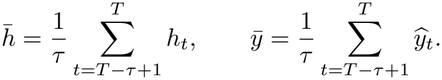

We used 30 recurrent steps and *τ* = 5. Input and readout matrices were initialized with orthogonal rows and held fixed throughout training, while the recurrent weight matrix was optimized. The initial weights were sampled as 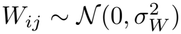, with 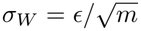 for *m* units with *ϵ* = 10^−4^. The bias *b* was optimized during training unless explicitly held fixed, as in the experiments comparing prescribed baseline levels.

Networks were trained by minimizing

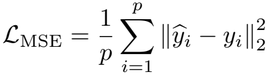

with Adam [29], where *p* is the number of training samples. We additionally included an activity regularization term to capture the metabolic cost associated with neural activity [30] and to place the network in the regime considered in our derivation of neural selectivity. The training objective therefore was

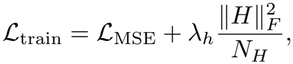

where *H* denotes the collection of hidden-unit activities over all training samples and, for recurrent networks, over all recurrent time steps; *N*_*H*_ is the total number of scalar entries in *H*. Gradients were clipped by their norm before each optimizer step. The learning rate, activity penalty, and training duration are specified below for each experiment. No input or recurrent noise was added in the simulations reported here.

### 9.2 Main Activation Pattern trees

The Main Activation Pattern (MAP) tree is a multi-output oblique decision tree fitted post hoc to approximate the routing performed by the neural network. Let 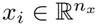 denote the input on trial *i* and *a*_*i*_ = (*a*_*i*1_, …, *a*_*im*_) ∈ {0, 1}^*m*^ its population activation pattern. The training data for the tree are therefore

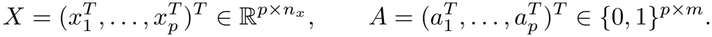

Unlike an ordinary classification tree, which predicts a single class, the MAP tree jointly predicts the *m* binary activation variables. Each internal node *v* applies an oblique split

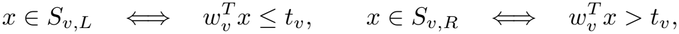

where *S*_*v*_ is the set of samples reaching the node, 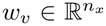 is a unit projection direction, and *t*_*v*_ is a scalar threshold. Thus, the leaves define a polyhedral partition of input space rather than a partition restricted to axis-aligned rectangles.

For a sample set *S*, the empirical activation probability of unit *j* is

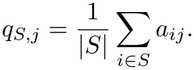

The node impurity used during fitting is the sample-weighted sum of the marginal binary entropies,

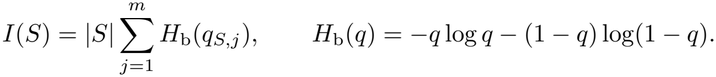

For a candidate split (*w, t*), the information gain is

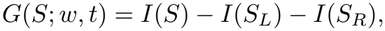

and the pair (*w, t*) producing the largest positive gain is retained. This marginal-entropy objective encourages each leaf to contain a reproducible population activation pattern while avoiding the need to treat every one of the potentially 2^*m*^ complete binary patterns as a separate class.

Candidate split directions were constructed at each node in three stages. First, we identified a candidate direction in input space along which the population activation pattern varied most strongly. This direction was estimated by ridge regression from the inputs to the binary activation patterns,

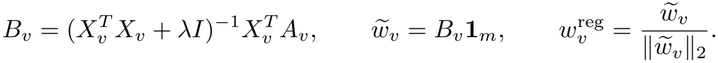

The ridge parameter was *λ* = 10^−6^. This direction, along with a set of normalized random directions, was included among the candidate split directions.

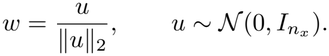

For each candidate direction *w*, the inputs were projected as *z*_*i*_ = *w*^*T*^ *x*_*i*_ and ordered along this one-dimensional axis. A possible split was evaluated between each pair of adjacent samples with different projected values, using their midpoint as the threshold. The selected split was the direction-threshold pair with the largest information gain,

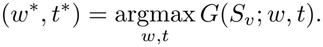

Finally, the selected direction was refined by testing normalized local perturbations of the form

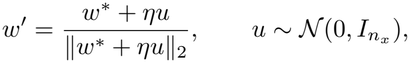

where *η* = 0.15 controls the refinement scale.

Trees were grown best-first. Every current leaf proposed its best admissible split, and the leaf with the greatest information gain was divided next. Consequently, a tree with *L* + 1 leaves was obtained from the *L*-leaf tree by adding the single split that most reduced total impurity. Growth stopped at the requested maximum number of leaves or when no positive-gain admissible split remained.

A leaf *l* stores the vector of empirical activation probabilities

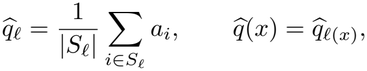

where *l*(*x*) is the leaf reached by input *x*. Tree fit was measured by the mean binary cross-entropy

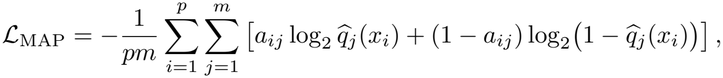

with probabilities bounded numerically away from zero and one. We considered two rules for converting the probability vector of leaf *l* into a binary activation pattern. Under a threshold rule,

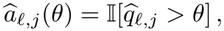

where *θ* is a fixed probability threshold. As a nonparametric alternative, the top-*k* rule preserves the rounded expected number of active neurons in each leaf. Specifically, we set

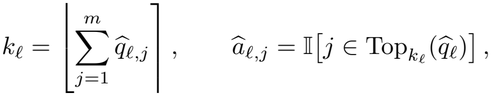

Where ⌊·⌉ denotes the nearest integer and Top_*k*_ (*q*) the indices of the *k* largest entries of *q*. Samples with the same resulting mask were treated as belonging to the same approximated activation region. The top-*k* rule provides a threshold-free alternative when a fixed cutoff is undesirable. Regional analyses excluded regions that did not contain enough samples or whose input matrix was numerically ill conditioned, as detailed below.

The conversion of continuous activity into the binary targets *A* differed between artificial ReLU networks and recorded neural data. In artificial networks, activity has an explicit mechanistic threshold: unit *j* was active precisely when its rectified activity was positive,

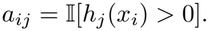

For feedforward networks, the activities of both hidden layers were concatenated before binarization, so that the MAP-tree target contained the complete multilayer activation pattern. For recurrent networks, activity was averaged over a terminal window of *τ* simulation steps,

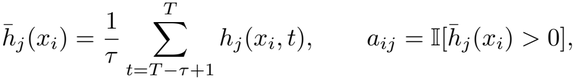

so that the tree described the stabilized end-of-trial computation rather than transient state changes. We used *τ* = 5 throughout.

Unlike artificial ReLU networks, recorded neural populations do not provide a known activation function from which binary activation states can be defined directly. We therefore introduced a plausible binarization rule: each neuron’s spike rate was standardized across trials, and responses above its mean firing rate were classified as active.

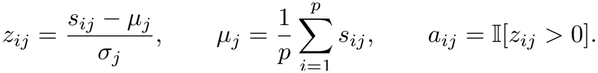

Thus, “active” denotes firing above that neuron’s trial-averaged rate, rather than firing above an absolute physiological threshold. This binary matrix was computed before the PCA-based processing used for the separate continuous selectivity analysis. After this initial binarization, artificial and recorded data followed the same MAP-tree fitting and binary-pattern prediction pipeline. Neurons with the same predicted activation pattern across leaves were grouped into the activation-defined subpopulations used for the selectivity analysis.

### 9.3 Feedforward nonlinear regression and regional least squares

The feedforward experiment in Fig. 3 used two hidden layers with 300 ReLU units each, two inputs, and one linear output. The input and output projection matrices *V* and *U* were initialized using orthogonal matrices and held fixed, while the hidden transformation *W* and the initially zero bias vectors *b*_1_ and *b*_2_ were trainable. The training set was a 40 × 40 regular grid over [−1.5, 1.5]^2^, with target

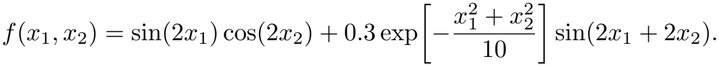

The network was trained for 1,000 optimizer steps using AdamW with learning rate 2 × 10^−3^, and activity regularizer *λ*_*h*_ = 10. Parameters and MAP trees were saved every ten optimizer steps. At each saved step, trees containing approximately logarithmically spaced numbers of leaves between 1 and 100 were fitted with a minimum of ten samples per leaf.

For every MAP-tree region, the network slope *A*_*r*_ and intercept *c*_*r*_ were computed from the effective weights associated with the predicted activation masks of the two hidden layers 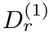 and 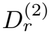:

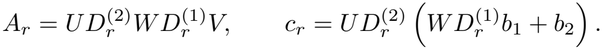

The corresponding affine map was written as *M*_*r*_ = (*c*_*r*_ | *A*_*r*_). Let *X*_*r*_ and *Y*_*r*_ contain the *p*_*r*_ inputs and targets assigned to region *r*, and define

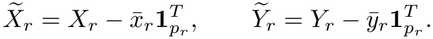

The regional affine least-squares slope and intercept were

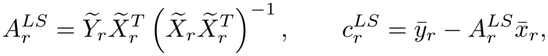

and the corresponding affine map was 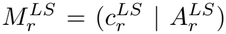. We quantified the deviation from the regional least-squares solution by Δ_*r*_, and the corresponding mean prediction error by ℰ_*r*_:

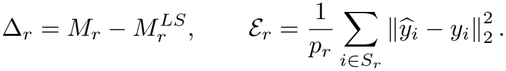

A region was retained when it contained at least ten samples and its centered input matrix had full input rank. To test the predicted square-root scaling, regional values from the largest fitted trees were pooled across training stages and fitted with

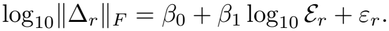

The slope, Pearson correlation coefficient, and two-sided *p* value are reported in Fig. 3.

### 9.4 Recurrent rule-switch task and fixed-point analysis

The context *c* ∈ {−1, 1} selected the relevant stimulus coordinate. The decision target was

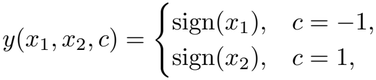

with zero assigned to the positive class.

The recurrent fixed-point experiment in Fig. 5 used 40 ReLU units, three inputs (*x*_1_, *x*_2_, *c*), and two outputs; the first reported the decision target defined above and the second reproduced the context. Stimulus coordinates were drawn uniformly from [−1, 1] and discretized into eight bins, and each training batch contained 120 trials. Networks were trained for 4,000 optimizer steps using AdamW with learning rate 2 × 10^−4^ and activity regularizer *λ*_*h*_ = 1. MAP trees with one to nine leaves were fitted to activation patterns averaged over the terminal window of length *τ* defined above.

Fixed-point predictions were evaluated for a trained network with baseline *b* = 5 using an eight-leaf MAP tree. For input *x*_*i*_ and predicted binary activation mask *a*_*i*_, the region-specific equilibrium was computed by solving the masked linear system

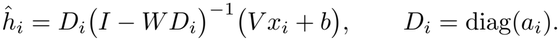

The global comparison omitted the mask and used the single linear solution obtained by setting *D*_*i*_ = *I*. Predictions were compared with the simulated activity at the final time step. Unit-wise accuracy was summarized by *R*^2^ = 1 − MSE*/* Var(*h*), and prediction errors were summarized across units by the natural logarithm of the mean squared error over trials. Principal-component displays were obtained by fitting PCA to the complete set of simulated trajectories and projecting both observed and predicted equilibria into the same space.

For the selectivity analysis, we used a separate rule-switch setup matching the sign and rotation experiments described below. The recurrent network contained 1,000 ReLU units, three inputs (*x*_1_, *x*_2_, *c*) and one linear output. Inputs were arranged on a homogeneous 16 × 16 grid over [−1, 1]^2^, repeated identically for both contexts. Each input was held constant throughout the recurrent sequence. The network was trained for 200 optimizer steps using AdamW with learning rate 10^−3^ and activity regularizer *λ*_*h*_ = 10. The task loss was the mean squared error over the final five recurrent steps.

### 9.5 Context-dependent sign task

The sign-task experiment used a recurrent network with 1,000 ReLU units, three inputs (*x*_1_, *x*_2_, *c*) and one linear output. The context *c* ∈ {−1, 1} selected between two nonlinear classification rules. The target was

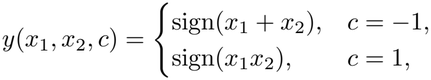

with zero assigned to the positive class. Inputs were arranged on a homogeneous 16 × 16 grid over [−1, 1]^2^, repeated identically in both contexts, yielding 512 task conditions. Each input was held constant throughout the recurrent sequence. The network was trained for 200 optimizer steps using AdamW with learning rate 10^−3^ and activity regularizer *λ*_*h*_ = 10. The task loss was the mean squared error over the final five recurrent steps.

### 9.6 Context-dependent rotation task

The rotation-task experiment used the same recurrent architecture, input grid, and training procedure. Here, context selected the orientation of a linear decision boundary. For context-dependent angles *θ*_−1_ = 0° and *θ*_+1_ = 60°, the target was

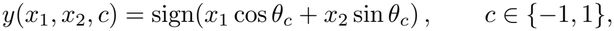

again assigning zero to the positive class. Thus, the relevant input direction was rotated by 60° between contexts while the sampled input distribution remained unchanged.

### 9.7 Baseline-dependent expressivity

To study how the operating point controls the number of activation regions, the rule-switch networks described above were trained from five independent initializations at baselines spanning the range used in the figures. For the systematic sweep, 40-unit networks were trained at 11 evenly spaced baselines between 0 and 5; additional high-baseline examples were trained to illustrate the staged learning regime. At each saved training stage, MAP trees were fitted and their cross-entropy was measured as a function of leaf number *L*. The improvement produced by adding one leaf was

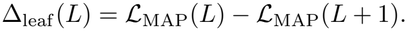

The effective number of regions was defined by the first addition for which Δ_leaf_ (*L*) *< δ*; the baseline sweep used *δ* = 0.03. Global one-plane and context-specific two-plane reference errors were obtained by ordinary least squares, either across all task conditions or separately within the two contexts. These reference errors identified plateaus at which a network behaved as one global affine transformation or as two context-specific transformations.

### 9.8 Finite-sample generalization experiment

The generalization experiment in Fig. 9 used recurrent networks with 300 ReLU units, two inputs and one output. The nonlinear radial target was

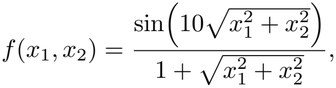

The training set comprised 61 points arranged on six concentric circles with radii evenly spaced between 0 and 1 and with 1, 4, 8, 12, 16, and 20 equally spaced angular samples, respectively. Networks were trained for 1,000 steps with AdamW, learning rate 10^−3^, no activity penalty. Eleven baseline values, *b* ∈ {0, 0.2, 0.4, 0.7, 1, 1.3, 1.6, 2, 2.5, 3, 4}, were tested with five independent network initializations per value.

Generalization error was evaluated on a 50 × 50 uniform grid *G* over [−0.8, 0.8]^2^ as

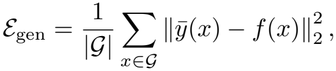

where 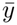 is the terminal-window average defined above. MAP trees with 1-120 leaves and at least five samples per leaf were fitted. A leaf *l* was designated trained when it contained at least one training input,

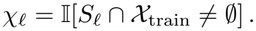

Errors were averaged separately over leaves with *χ*_*l*_ = 1 and *χ*_*l*_ = 0. The paired comparison in Fig. 9 used a two-sided Wilcoxon signed-rank test across matched network fits (*W* = 29, *p* = 3.16 × 10^−9^). Curves and shaded bands show the mean and mean ± one standard error across the five initializations.

### 9.9 Empirical data pre-processing

#### 9.9.1 Reinert dataset

Recordings from different animals and sessions were combined into a pseudo-population. Trials were divided into eight conditions defined by task context, stimulus category, and behavioral outcome (hit, miss, correct rejection, or false alarm). For each condition, the number of pseudo-trials was set to the mean number of available trials across recordings. Trials were then sampled with replacement within each recording and condition, and the resulting activity vectors were concatenated across recordings.

Let *H* ∈ ℝ^*p*×*n*^ denote the pseudo-trial-by-neuron spike-rate matrix. Each neuron’s activity was standardized across pseudo-trials as

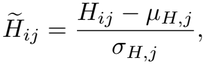

where *µ*_*H,j*_ and *σ*_*H,j*_ are the mean and standard deviation of neuron *j* across pseudo-trials. The binary activation-pattern matrix *A* ∈ {0, 1}^*p*×*n*^ was then defined before PCA denoising by

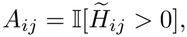

where *A*_*ij*_ = 1 indicates that neuron *j* was active on pseudo-trial *i*. For the continuous selectivity analysis, 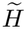 was transformed onto its first ten principal components and inverse transformed in the original neuron space, thereby retaining the dominant shared population activity modes.

Let *o*_*i*_, *s*_*i*_ ∈ {−1, +1} denote the orientation and spatial-frequency categories on pseudo-trial *i*, respectively, and let *c*_*i*_ denote its task context. The input and target vectors were

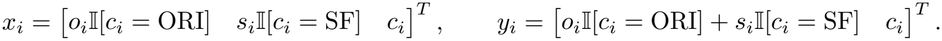

The first two input coordinates describe context-gated orientation and spatial-frequency evidence. The first target coordinate encodes the context-dependent NoGo/Go action as −1*/*+1, and the second encodes context. The columns of the resulting input and target matrices *X* and *Y* were standardized across pseudo-trials as

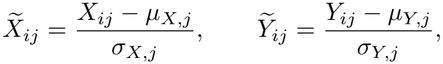

where *µ*_*X,j*_ and *σ*_*X,j*_, respectively *µ*_*Y,j*_ and *σ*_*Y,j*_, are the mean and standard deviation of column *j* across pseudo-trials.

#### 9.9.2 Hajnal dataset

Only correct trials from the complex task blocks were retained. Visual orientation (45° or 135°), auditory frequency (5 or 10 kHz), and task context (auditory or visual) were encoded as binary variables in {−1, +1}. ACC and V1 recordings were pooled for the analysis reported here. For each neuron and trial, the response was defined as the mean firing rate during the 1.5-s interval following trial onset.

The three binary task variables defined eight experimental conditions. For each condition, the number of pseudo-trials was set to the mean number of available trials across recordings. Each pseudo-trial was constructed by independently sampling, with replacement, one response from the corresponding condition for every neuron. The sampled responses were concatenated across neurons to form the pseudo-population activity matrix *H*.

Each neuron’s activity was standardized across pseudo-trials as

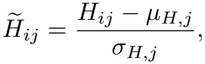

where *µ*_*H,j*_ and *σ*_*H,j*_ are the mean and standard deviation of neuron *j* across pseudo-trials. The binary activation-pattern matrix *A* ∈ {0, 1}^*p*×*n*^ was defined before PCA denoising by

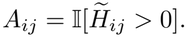

For the continuous selectivity analysis, 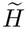 was transformed onto its first ten principal components and inverse transformed in the original neuron space, thereby retaining the dominant shared population activity modes.

Let *o*_*i*_, *f*_*i*_ ∈ {−1, +1} denote visual orientation and auditory frequency, respectively, and let *c*_*i*_ denote task context. The input and target vectors were constructed in the same way as for the Reinert dataset,

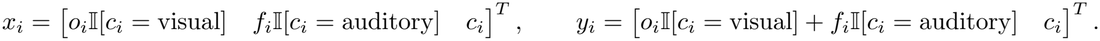

Thus, the first target coordinate encoded the correct NoGo/Go action as −1*/* + 1, and the second encoded context. The columns of the resulting input and target matrices *X* and *Y* were standardized across pseudo-trials as

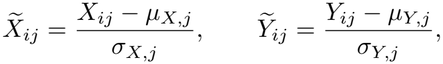

where *µ*_*X,j*_ and *σ*_*X,j*_, respectively *µ*_*Y,j*_ and *σ*_*Y,j*_, are the mean and standard deviation of column *j* across pseudo-trials.

### 9.10 Neural-data selectivity analysis

The selectivity coefficients were estimated from the standardized matrices by ordinary least squares,

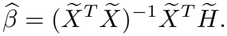

Each column of 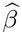 contains the selectivity coefficients of one neuron with respect to the task variables represented in 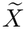.

A MAP tree was fitted to predict the binary activation-pattern matrix *A* from the standardized task variables 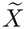, with at least ten pseudo-trials per leaf. The covariance analyses used four-leaf trees throughout, except for the simulated sign task, for which a seven-leaf tree was used. Leaf probabilities were converted into binary patterns using the threshold rule, with *θ* = 0.1 for the three simulated tasks, *θ* = 0.2 for the Reinert dataset, and *θ* = 0.3 for the Hajnal dataset. For each dataset, the threshold was selected to maximize the diversity of the resulting binary activation patterns. Neurons were grouped into subpopulations according to the set of tree regions in which they were predicted to be active. Subpopulations with zero predicted covariance or containing less than 5% of the recorded neurons were excluded.

For a retained subpopulation *s* containing *n*_*s*_ neurons, let *β*_*j*_ ∈ ℝ^3^ be the selectivity vector of neuron *j*. Its measured selectivity geometry was

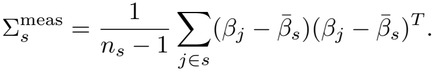

To predict this geometry from task statistics, the standardized input variables 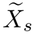 from leaves in which the subpopulation was active were regressed onto the corresponding standardized task outputs 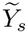,

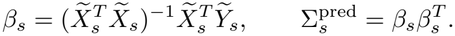

Agreement was quantified by the cosine similarity between the vectors containing the unique entries of 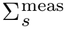 and 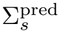. The corresponding similarity and rotation-test *p* value are displayed for each subpopulation in Fig. 6.

### 9.11 Statistical reporting

For the selectivity-geometry analysis, significance was assessed using Haar-distributed rotations. At each permutation *k*, a *d* × *d* matrix *G*_*k*_ was generated with independent entries (*G*_*k*_)_*ij*_ ~ N (0, 1), where *d* is the number of selectivity dimensions, and decomposed as *G*_*k*_ = *Q*_*k*_*R*_*k*_ by QR decomposition. The resulting matrix *Q*_*k*_ satisfies 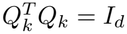 and defines a random rotation with no preferred orientation [31]. The predicted covariance matrix of every retained subpopulation was then rotated as 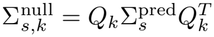. The same rotation was applied to all subpopulations within a permutation. If *z*_*s*_ denotes the observed cosine similarity and 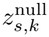 its value after rotation, the population-dependent one-sided *p* value was

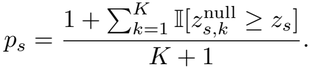

The aggregate statistic was the mean cosine similarity across the fixed set *S* of retained subpopulations, 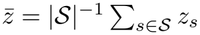. Its null distribution was obtained by averaging across the same subpopulations after each common rotation, 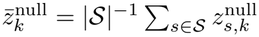, and its *p* value was computed using the same upper-tail formula. All analyses used *K* = 1,000 rotations.

## 10 Supplementary Figures

**Supplementary Fig. 1.**
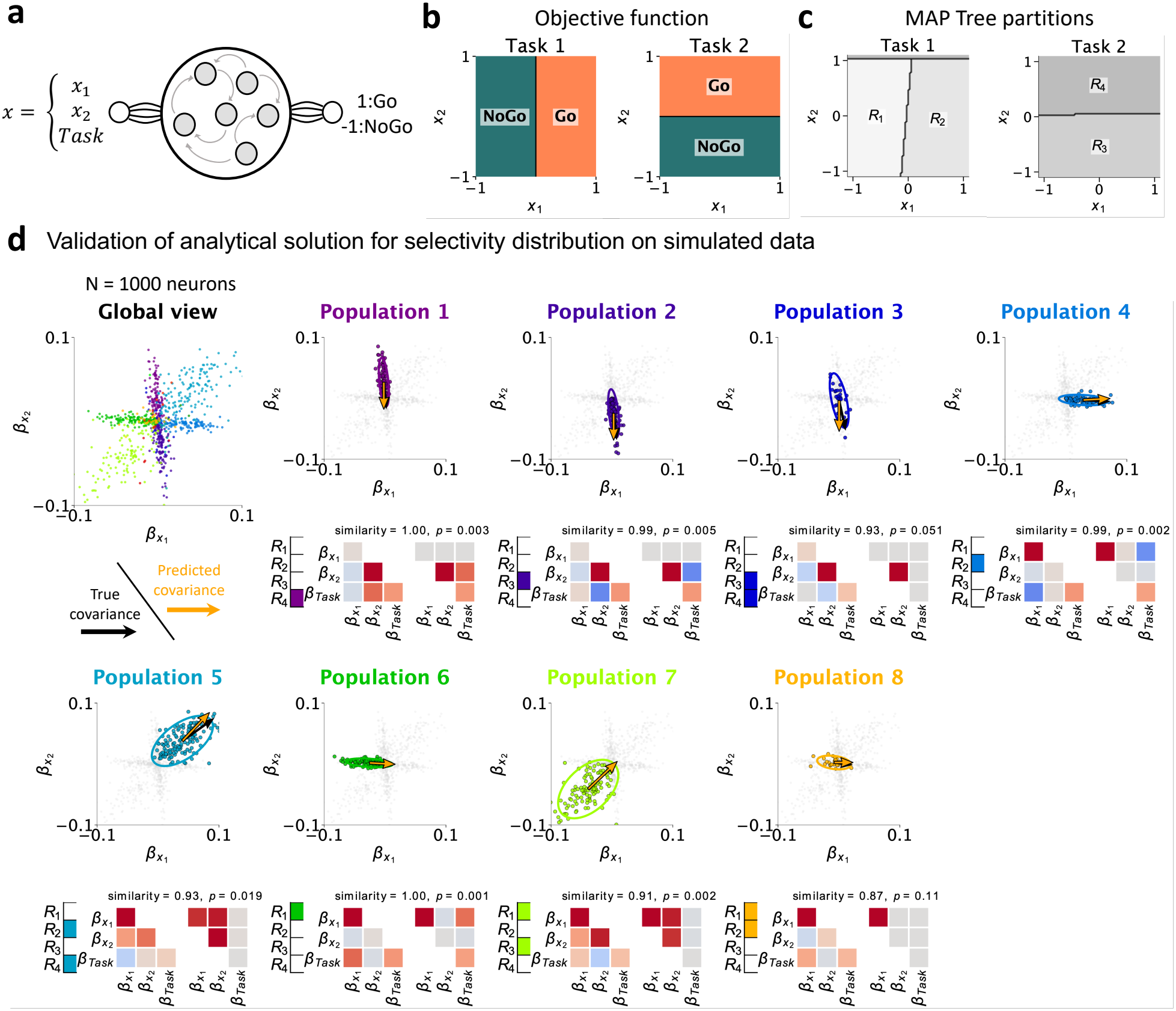
Task statistics predict the selectivity geometry of activation-defined neural subpopulations in the simulated rule-switch task. **a**, Recurrent-network architecture. The network received two stimulus features, *x*_1_ and *x*_2_, together with a task cue, and produced a binary go/no-go response. **b**, Objective functions in the two task contexts. The response depended on the sign of *x*_1_ in Task 1 and on the sign of *x*_2_ in Task 2; black lines indicate the category boundaries. **c**, Partition of the input space by a four-leaf MAP tree fitted to network activity. Gray levels distinguish the regions *R*_1_-*R*_4_ in each task context. **d**, Validation of the predicted selectivity geometry in 1,000 simulated neurons. The global view shows all neurons in selectivity space, color-coded by activation-defined subpopulation, and the detailed plots show each subpopulation separately. Colored points identify the selected subpopulation against the remaining neurons in gray, and ellipses represent measured covariance. Black and orange arrows indicate the principal directions of the measured and theoretically predicted covariance, respectively. The adjacent region indicators identify the MAP-tree regions in which each subpopulation was active. Covariance matrices compare the measured lower triangle with the theoretically predicted upper triangle across *x*_1_, *x*_2_, and task selectivities. Matrix entries are rescaled coefficients ranging from −1 (blue) to +1 (red). The displayed similarity is the cosine similarity between the unique entries of the measured and predicted covariance matrices; the associated *p* value is obtained from the rotation-based permutation test described in Methods.

**Supplementary Fig. 2.**
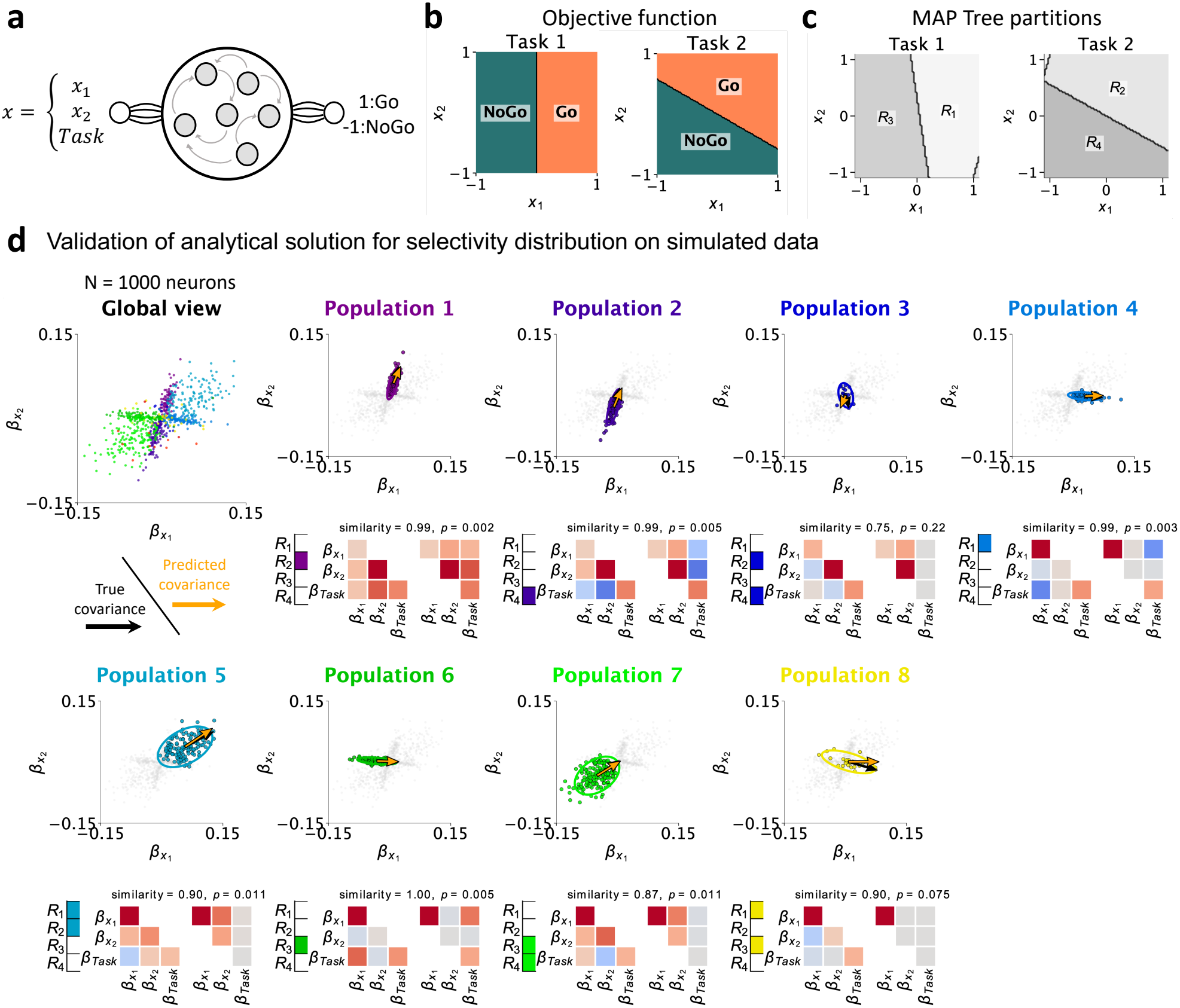
Task statistics predict the selectivity geometry of activation-defined neural subpopulations in the simulated context-dependent rotation task. **a**, Recurrent-network architecture. The network received two stimulus features, *x*_1_ and *x*_2_, together with a task cue, and produced a binary go/no-go response. **b**, Objective functions in the two task contexts. The decision boundary was rotated from 0° in Task 1 to 60° in Task 2; black lines indicate the category boundaries. **c**, Partition of the input space by a four-leaf MAP tree fitted to network activity. Gray levels distinguish the regions *R*_1_-*R*_4_ in each task context. **d**, Validation of the predicted selectivity geometry in 1,000 simulated neurons. The global view shows all neurons in selectivity space, color-coded by activation-defined subpopulation, and the detailed plots show each subpopulation separately. Colored points identify the selected subpopulation against the remaining neurons in gray, and ellipses represent measured covariance. Black and orange arrows indicate the principal directions of the measured and theoretically predicted covariance, respectively. The adjacent region indicators identify the MAP-tree regions in which each subpopulation was active. Covariance matrices compare the measured lower triangle with the theoretically predicted upper triangle across *x*_1_, *x*_2_, and task selectivities. Matrix entries are rescaled coefficients ranging from −1 (blue) to +1 (red). The displayed similarity is the cosine similarity between the unique entries of the measured and predicted covariance matrices; the associated *p* value is obtained from the rotation-based permutation test described in Methods.

**Supplementary Fig. 3.**
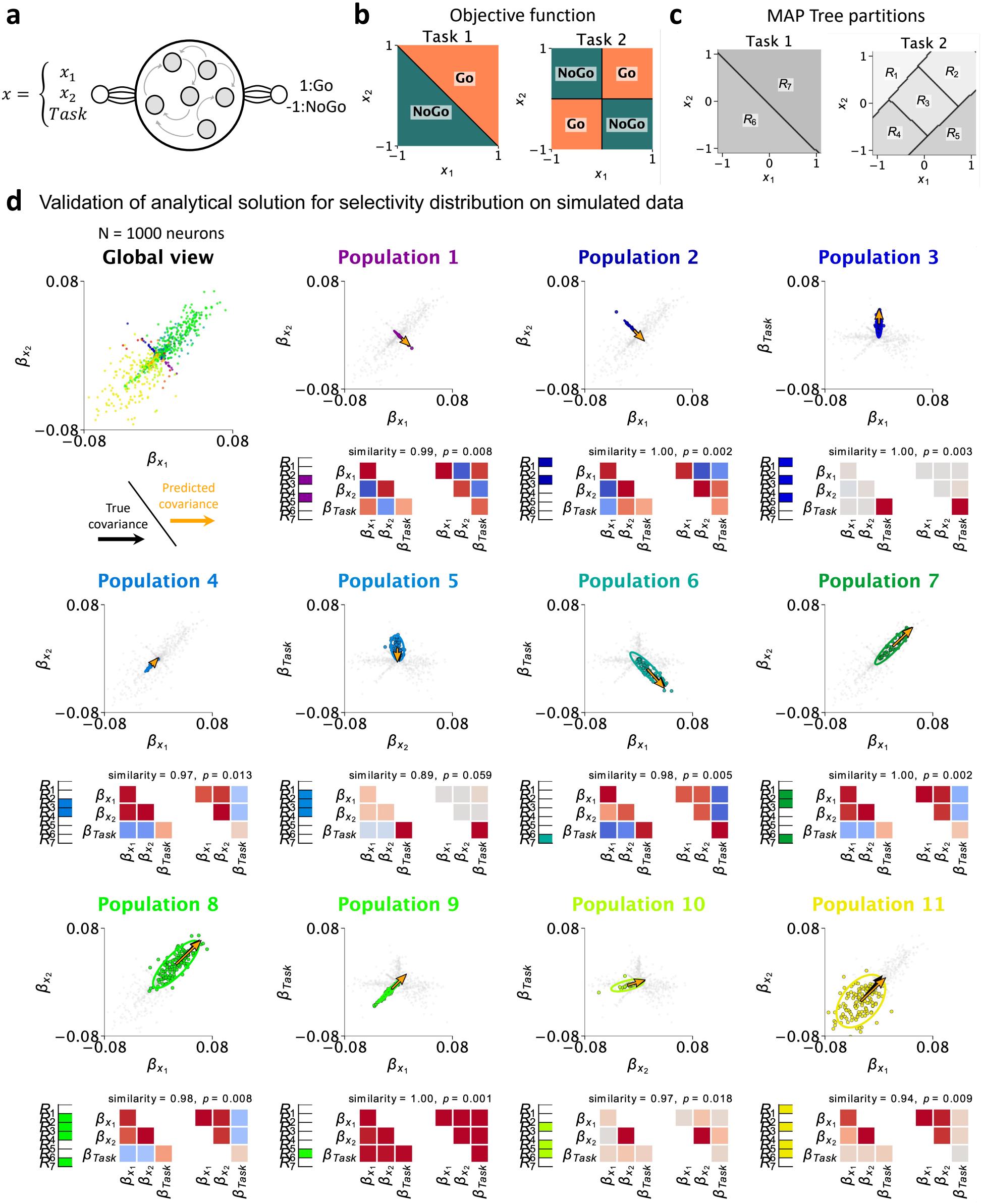
Task statistics predict the selectivity geometry of activation-defined neural subpopulations in the simulated context-dependent sign task. **a**, Recurrent-network architecture. The network received two stimulus features, *x*_1_ and *x*_2_, together with a task cue, and produced a binary go/no-go response. **b**, Objective functions in the two task contexts. The response was determined by the sign of *x*_1_ + *x*_2_ in Task 1 and by the sign of *x*_1_*x*_2_ in Task 2; black lines indicate the category boundaries. **c**, Partition of the input space by a seven-leaf MAP tree fitted to network activity. Gray levels distinguish the regions *R*_1_-*R*_7_ in each task context. **d**, Validation of the predicted selectivity geometry in 1,000 simulated neurons. The global view shows all neurons in selectivity space, color-coded by activation-defined subpopulation, and the detailed plots show each subpopulation separately. Colored points identify the selected subpopulation against the remaining neurons in gray, and ellipses represent measured covariance. Black and orange arrows indicate the principal directions of the measured and theoretically predicted covariance, respectively. The adjacent region indicators identify the MAP-tree regions in which each subpopulation was active. Covariance matrices compare the measured lower triangle with the theoretically predicted upper triangle across *x*_1_, *x*_2_, and task selectivities. Matrix entries are rescaled coefficients ranging from −1 (blue) to +1 (red). The displayed similarity is the cosine similarity between the unique entries of the measured and predicted covariance matrices; the associated *p* value is obtained from the rotation-based permutation test described in Methods.

**Supplementary Fig. 4.**
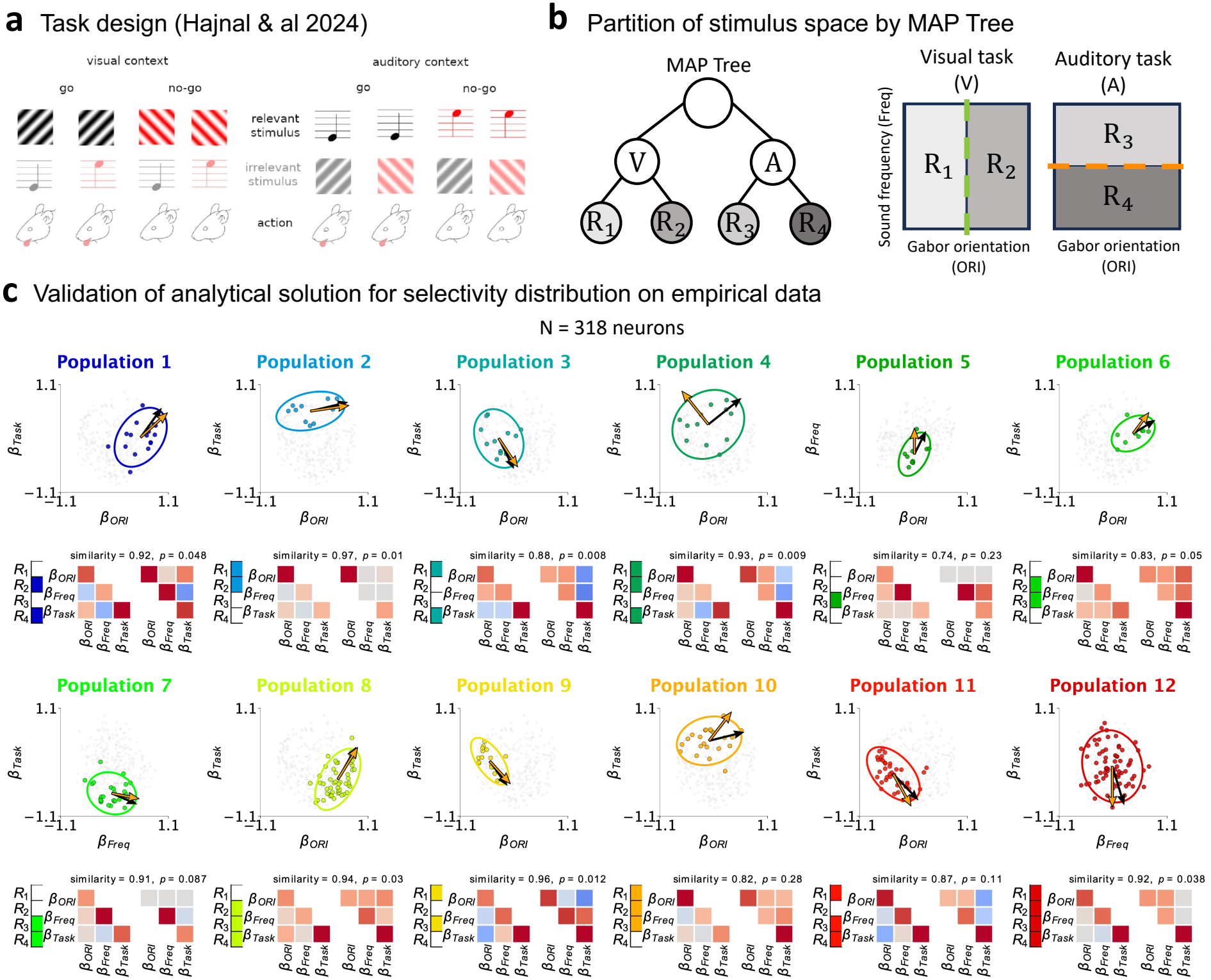
Task statistics predict the selectivity geometry of activation-defined neural subpopulations in the Hajnal dataset. **a**, Task design adapted from Hajnal et al. (2024). Mice integrated visual orientation and auditory frequency according to the current context to produce a go or no-go response. **b**, Partition of the joint stimulus space by a MAP tree fitted to population activity. The first split separates the visual (V) and auditory (A) contexts, and subsequent splits along the stimulus dimensions define four activation regions (*R*_1_-*R*_4_), shown in the corresponding stimulus spaces. Green and orange dashed lines indicate the category boundaries. **c**, Validation of the predicted selectivity geometry in 318 recorded neurons. The twelve detailed plots show activation-defined subpopulations corresponding to distinct combinations of MAP-tree regions. In each plot, colored points highlight the selected subpopulation against the remaining neurons in gray, and the ellipse represents its measured covariance in the displayed pair of selectivity dimensions. Black and orange arrows indicate the principal directions of the measured and theoretically predicted covariance, respectively. The adjacent *R*_1_-*R*_4_ indicators identify the activation regions defining each subpopulation. Covariance matrices compare the measured lower triangle with the theoretically predicted upper triangle across orientation, frequency, and task selectivities. Matrix entries are rescaled coefficients ranging from −1 (blue) to +1 (red). The displayed similarity is the cosine similarity between the unique entries of the measured and predicted covariance matrices; the associated *p* value is obtained from the rotation-based permutation test described in Methods.

## References

1. Saxe, A. M., McClelland, J. L. & Ganguli, S. Exact solutions to the nonlinear dynamics of learning in deep linear neural networks. 2014. doi: 10.48550/arXiv.1312.6120.

2. Saxe, A. M., McClelland, J. L. & Ganguli, S. A mathematical theory of semantic development in deep neural networks. Proceedings of the National Academy of Sciences 116 (2019), pp. 11537–11546. doi: 10.1073/pnas.1820226116.

3. Chu, L., Hu, X., Hu, J., Wang, L. & Pei, J. Exact and Consistent Interpretation for Piecewise Linear Neural Networks: A Closed Form Solution. Proceedings of the 24th ACM SIGKDD International Conference on Knowledge Discovery & Data Mining. London United Kingdom: ACM, 2018, pp. 1244–1253. doi: 10.1145/3219819.3220063.

4. Balestriero, R. & baraniuk. A Spline Theory of Deep Learning. Proceedings of the 35th International Conference on Machine Learning. PMLR, 2018, pp. 374–383.

5. Montúfar, G., Pascanu, R., Cho, K. & Bengio, Y. On the Number of Linear Regions of Deep Neural Networks. 2014. doi: 10.48550/arXiv.1402.1869.

6. Serra, T., Tjandraatmadja, C. & Ramalingam, S. Bounding and Counting Linear Regions of Deep Neural Networks. Proceedings of the 35th International Conference on Machine Learning. PMLR, 2018, pp. 4558–4566.

7. Hanin, B. & Rolnick, D. Complexity of Linear Regions in Deep Networks. Proceedings of the 36th International Conference on Machine Learning. PMLR, 2019, pp. 2596–2604.

8. Goujon, A., Etemadi, A. & Unser, M. On the Number of Regions of Piecewise Linear Neural Networks. 2023. doi: 10.48550/arXiv.2206.08615.

9. Rigotti, M. et al. The importance of mixed selectivity in complex cognitive tasks. Nature 497 (2013), pp. 585–590. doi: 10.1038/nature12160.

10. Mante, V., Sussillo, D., Shenoy, K. V. & Newsome, W. T. Context-dependent computation by recurrent dynamics in prefrontal cortex. Nature 503 (2013), pp. 78–84. doi: 10.1038/nature12742.

11. Yang, G. R., Joglekar, M. R., Song, H. F., Newsome, W. T. & Wang, X.-J. Task representations in neural networks trained to perform many cognitive tasks. Nature Neuroscience 22 (2019), pp. 297–306. doi: 10.1038/s41593-018-0310-2.

12. Reinert, S., Hübener, M., Bonhoeffer, T. & Goltstein, P. M. Mouse prefrontal cortex represents learned rules for categorization. Nature 593 (2021), pp. 411–417. doi: 10.1038/s41586-021-03452-z.

13. Dubreuil, A., Valente, A., Beiran, M., Mastrogiuseppe, F. & Ostojic, S. The role of population structure in computations through neural dynamics. Nature neuroscience 25 (2022), pp. 783–794. doi: 10.1038/s41593-022-01088-4.

14. Barbosa, J. et al. Early selection of task-relevant features through population gating. Nature Communications 14 (2023), p. 6837. doi: 10.1038/s41467-023-42519-5.

15. Tissot, H., Boucher, J., Reinert, S., Goltstein, P. M. & Boubenec, Y. Sensorimotor remapping drives task specialization in prefrontal cortex. Nature Communications 17 (2026), p. 9904. doi: 10.1038/s41467-026-76104-3.

16. Hajnal, M. A. et al. Shifts in attention drive context-dependent subspace encoding in anterior cingulate cortex in mice during decision making. Nature Communications 15 (2024), p. 5559. doi: 10.1038/s41467-024-49845-2.

17. Nguyen, D. T., Kasmarik, K. E. & Abbass, H. A. Towards Interpretable ANNs: An Exact Transformation to Multi-Class Multivariate Decision Trees. 2021. doi: 10.48550/arXiv.2003.04675.

18. Aytekin, C. Neural Networks are Decision Trees. 2022. doi: 10.48550/arXiv.2210.05189.

19. Bernardi, S. et al. The Geometry of Abstraction in the Hippocampus and Prefrontal Cortex. Cell 183 (2020), 954–967.e21. doi: 10.1016/j.cell.2020.09.031.

20. Hahnloser, R. H. R., Sarpeshkar, R., Mahowald, M. A., Douglas, R. J. & Seung, H. S. Digital selection and analogue amplification coexist in a cortex-inspired silicon circuit. Nature 405 (2000), pp. 947–951. doi: 10.1038/35016072.

21. Salinas, E & Abbott, L. Transfer of coded information from sensory to motor networks. The Journal of Neuroscience 15 (1995), pp. 6461–6474. doi: 10.1523/JNEUROSCI.15-10-06461.1995.

22. Chance, F. S., Abbott, L. & Reyes, A. D. Gain Modulation from Background Synaptic Input. Neuron 35 (2002), pp. 773–782. doi: 10.1016/S0896-6273(02)00820-6.

23. Arieli, A., Sterkin, A., Grinvald, A. & Aertsen, A. Dynamics of Ongoing Activity: Explanation of the Large Variability in Evoked Cortical Responses. Science 273 (1996), pp. 1868–1871. doi: 10.1126/science.273.5283.1868.

24. Churchland, M. M. et al. Stimulus onset quenches neural variability: a widespread cortical phenomenon. Nature Neuroscience 13 (2010), pp. 369–378. doi: 10.1038/nn.2501.

25. Sussillo, D. & Barak, O. Opening the Black Box: Low-Dimensional Dynamics in High-Dimensional Re-current Neural Networks. Neural Computation 25 (2013), pp. 626–649. doi: 10.1162/NECO_a_00409.

26. Chaisangmongkon, W., Swaminathan, S. K., Freedman, D. J. & Wang, X.-J. Computing by Robust Transience: How the Fronto-Parietal Network Performs Sequential, Category-Based Decisions. Neuron 93 (2017), 1504–1517.e4. doi: 10.1016/j.neuron.2017.03.002.

27. Driscoll, L. N., Pettit, N. L., Minderer, M., Chettih, S. N. & Harvey, C. D. Dynamic Reorganization of Neuronal Activity Patterns in Parietal Cortex. Cell 170 (2017), 986–999.e16. doi: 10.1016/j.cell.2017.07.021.

28. Kirkpatrick, J. et al. Overcoming catastrophic forgetting in neural networks. Proceedings of the National Academy of Sciences 114 (2017), pp. 3521–3526. doi: 10.1073/pnas.1611835114.

29. Kingma, D. P. & Ba, J. Adam: A Method for Stochastic Optimization. 2017. doi: 10.48550/arXiv.1412.6980.

30. Attwell, D. & Laughlin, S. B. An Energy Budget for Signaling in the Grey Matter of the Brain. Journal of Cerebral Blood Flow & Metabolism 21 (2001), pp. 1133–1145. doi: 10.1097/00004647-200110000-00001.

31. Mezzadri, F. How to generate random matrices from the classical compact groups. 2007. doi: 10.48550/arXiv.math-ph/0609050.

